# Anchored for action: a dual role for integrin β1 in mast cell perivascular positioning and vasoactivity to license leukocyte recruitment

**DOI:** 10.64898/2026.05.07.723399

**Authors:** Aaron Hoffmann, Sebastian Drube, Roland Immler, Konstantinos Katsoulis-Dimitriou, Jan Dudeck, Kathleen Baumgart, Claudia Küchler, Tobias Franz, Stephan Fricke, Sascha Kahlfuß, Markus Sperandio, Anne Dudeck

**Affiliations:** Institute for Clinical Immunology and Cell Therapeutics, Medical Faculty, Otto-von-Guericke University Magdeburg, Germany; Institute for Immunology, University Hospital Jena, Jena, Germany; Institute for Cardiovascular Physiology and Pathophysiology, Biomedical Center (BMC), Ludwig-Maximilians-University of Munich, Planegg-Martinsried, Germany; Multi-parametric Bioimaging and Cytometry platform, Medical Faculty, Otto-von-Guericke University Magdeburg, Germany; Health Campus Immunology, Infectiology and Inflammation, Otto-von-Guericke-University, Magdeburg, Germany; Fraunhofer Institute for Cell Therapy and Immunology, Leipzig, Germany

**Keywords:** mast cell, integrin β1, intravascular degranulation, skin inflammation, vasoactivity

## Abstract

Mast cells (MCs) are tissue-resident sentinels of the innate immune system that play pivotal roles in host defense and inflammation. Perivascular MCs exert a particularly strong influence on the onset and dynamics of inflammation through the rapid, directional release of proinflammatory mediators into the circulation. Yet, the mechanisms governing their attachment to the vessel wall – a prerequisite for intravascular degranulation – remain poorly defined. Using a conditional knockout of integrin β1 (Itgb1) in MCs, we investigated how perivascular positioning, degranulation, and vasoactive function contribute to inflammatory responses. *In vivo* imaging revealed that Itgb1 is essential for positioning MCs within the perivascular niche, particularly around arterioles. The absence of Itgb1 markedly reduced directional MC degranulation into blood vessels during skin inflammation. *In vitro*, Itgb1-deficient MCs displayed impaired degranulation kinetics together with altered SHIP1/PI3K-AKT signaling and calcium influx upon P2X7 ligation by ATP. During contact hypersensitivity, mice lacking Itgb1 in MCs exhibited strongly diminished ear swelling and reduced recruitment of multiple leukocyte subsets. Mechanistically, disordered MC positioning and attenuated degranulation impaired endothelial activation, resulting in decreased leukocyte adhesion and extravasation. These findings uncover a dual role for Itgb1 in regulating MC responsiveness and pro-inflammatory vasoactive function, establishing Itgb1-mediated perivascular MC positioning as a key prerequisite for effective leukocyte recruitment.

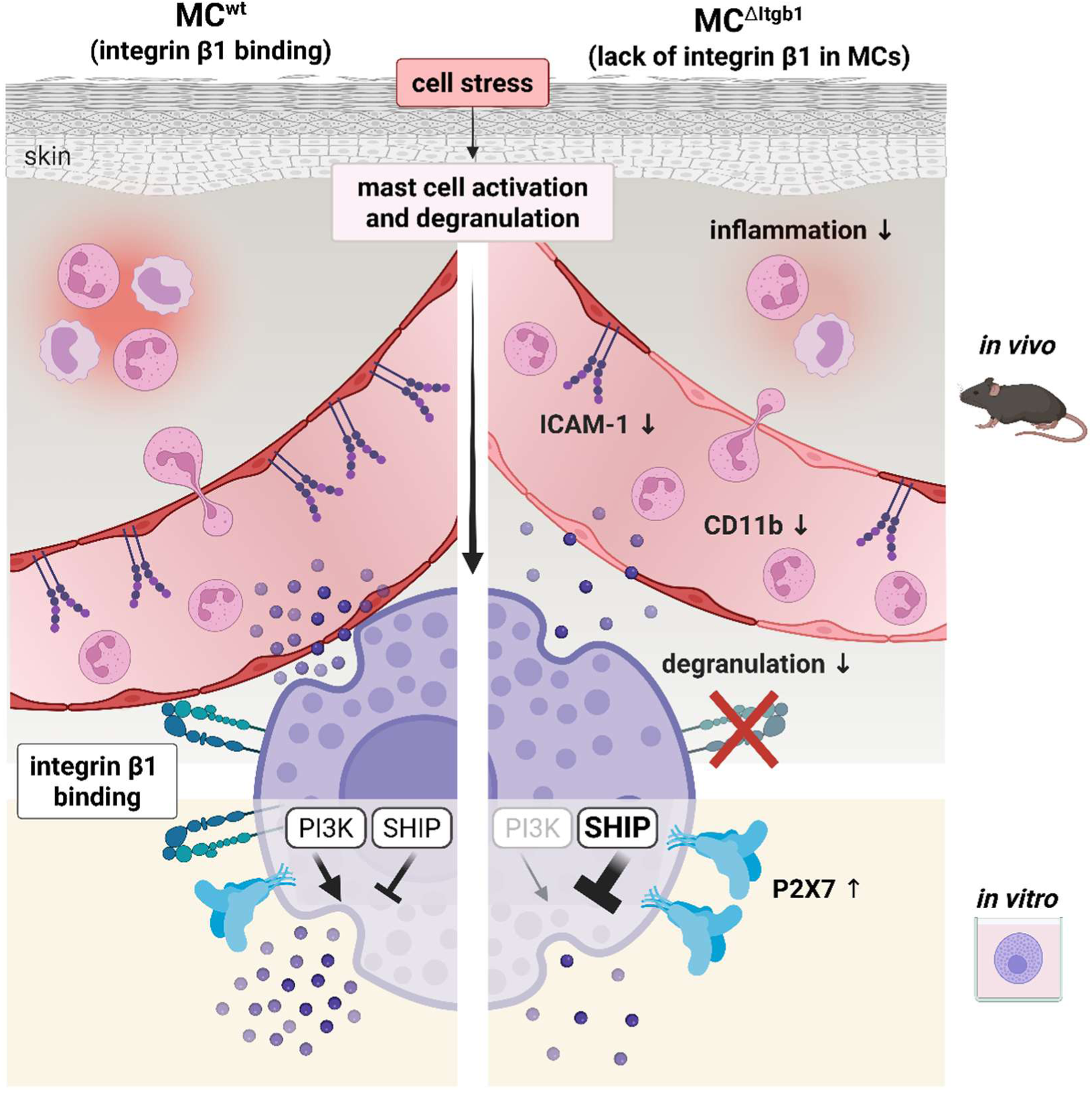

## INTRODUCTION

Nearly 20% of the general population suffer from allergic contact dermatitis (ACD), a type IV allergic response, which has severe negative socio-economic consequences for our society (1–3). The mouse model of contact hypersensitivity (CHS) has highlighted the pivotal role of mast cells (MCs) in ACD, by both pro-and anti-inflammatory characteristics (4–8). Beyond their well characterized effector functions in immediate type allergic responses and venom degradation, MCs critically contribute to host defense and leukocyte recruitment (9,10). Activation of the FcεR/IgE complex or a variety of other exogenous and endogenous noxes instigates the prompt release of pre-formed mediators, including histamine, tumor necrosis factor (TNF) and various other cytokines, which are embedded in secretory granules (9,11,12). In CHS, sensitization by the organic hapten 1-fluoro-2,4-dinitrobenzene (DNFB) induces the release of the alarmins adenosine triphosphate (ATP) and interleukin (IL) 33 by stressed keratinocytes, leading to MC degranulation and priming of hapten-specific T cells. Subsequent contact with the hapten (challenge) initiates a T cell-driven delayed type hypersensitivity response with local inflammation, MC activation, increased vascular permeability and neutrophil influx (5,8,13). Notably, perivascular MCs critically impact on the onset and kinetics of inflammation. Via intraluminal sheets, MCs facilitate the directional release of TNF into the bloodstream, thereby priming circulating neutrophils as a precondition for blood vessel extravasation and skin infiltration (14). Thus, local MC activation may lead to systemic distribution of inflammatory mediators via the blood circulation, with the risk of potentially life-threatening anaphylactic shock syndromes or cytokine-storm symptoms in humans. However, the mechanisms underlying MC attachment to the vessel wall, as a prerequisite for the intravascular degranulation and subsequent vasoactive and leukocytes recruiting functions, are not yet fully understood.

For cell-cell communication and interaction with other cells or the extracellular matrix (ECM), MCs express many adhesion molecules, including integrins, selectins, and cadherins (15). In contrast to other leukocytes, MC migration, adhesion and perivascular blood vessel alignment is fully dependent on integrins, highlighting their key role for MC positioning in homeostasis (16). The integrin family comprises 18 α subunits and 8 β subunits, resulting in 24 distinct heterodimers (17). Many of these subunits are expressed by MCs, with notable expression of αV, β1 and β3 in mature MCs (15,16). Ligation of integrin receptors mediates MC motility, survival, effector functions and apoptosis (17). In particular, integrin β1 (Itgb1) was recently shown to be critical for MC-blood vessel alignment and immune surveillance of blood vessel content (18). However, the mechanistic effect of Itgb1 binding on MC activation, degranulation and, thus, inflammation-driver functions remains elusive. A deeper understanding of these processes could potentially help to develop targeted, MC-specific treatment to prevent severe effects upon MC-activation.

In this study, we employed a Mcpt5-Cre-mediated conditional gene inactivation of Itgb1 (MC^ΔItgb1^) to deplete the group of Itgb1 receptors in connective tissue type MCs (CTMCs) that are involved in the binding to fibronectin, collagen, laminin and VCAM-1 (15). Here, we demonstrate that Itgb1 is essential for the blood vessel positioning and elongated morphology of perivascular MCs, but more importantly is critical for MC activation and degranulation *in vivo* and *in vitro*. Non-adherent MCs showed an aberrant degranulation efficiency in the absence of Itgb1 *in vitro*, accompanied by altered signaling patterns and calcium influx upon ATP stimulation, underlining cell intrinsic Itgb1 effects, independent of ECM binding. MC^ΔItgb1^ mice showed a diminished directional intravascular degranulation capacity. Interestingly, the aberrant perivascular alignment and reduced degranulation of MCs in the absence of Itgb1 resulted in abrogated activation of the blood endothelium, along with impaired neutrophil recruitment and extravasation *in vivo*. This, in turn, links back to the markedly diminished leukocyte infiltration in CHS. Thus, our findings reveal a dual role for Itgb1 in the responsiveness of perivascular MCs and their pro-inflammatory vasoactive effects on endothelial cell activation, which are crucial for leukocyte recruitment in CHS.

## RESULTS

We have previously demonstrated that perivascular MCs play a central role in inflammation by the directional degranulation into the blood stream (14). Following this observation, we aimed to identify the mechanisms of vessel alignment and penetration, as well as the morphological and functional characteristics of perivascular MCs in contrast to non-vessel associated interstitial MCs.

### Blood vessel alignment and elongated morphology of perivascular MCs requires integrin β1

To investigate the relevance of the adhesion molecule integrin β1 (Itgb1) in blood vessel attachment and alignment of perivascular MCs, we crossed the Itgb1^fl/fl^ mouse line (19) to Mcpt5-Cre mice to generate a conditional Itgb1 knock-out in Cre-expressing connective-tissue type MCs (CTMCs, referred to as MC^ΔItgb1^, Fig. 1A). In line with another publication (18), the majority of ear skin and primary isolated and differentiated peritoneal cell-derived MCs (PCMCs) revealed a high expression of Itgb1 in MC^wt^, while no Itgb1 expression was detected in MC^ΔItgb1^ (Fig. 1A-C). Further, Itgb1 was proved to be expressed in the open and active conformation in MC^wt^ (Fig. 1D-E). Subsequent analyses revealed that other leukocyte subsets remained unaltered with respect to their expression of Itgb1 and their frequency (Fig. S1A, B). Mature MCs express several alpha chains pairing with Itgb1, such as integrin α2, α3, α4, α5, and also integrin β3, but not β2 (15). However, integrin β3 expression was not altered in MC^ΔItgb1^ (Fig. S1C), underlining the suitability of this mouse line for the functional investigation of Itgb1 in CTMCs. As previously described (14,18) and confirmed by fluorescence microscopy, perivascular ear skin MCs from MC^wt^ mice are elongated and in close proximity to the blood vessels. In contrast, Itgb1-deficient MCs showed a drastically altered morphology (Fig. 1F). While the total MC number was unaffected by the Itgb1-ko (Fig. 1G), cell-based parameter quantification confirmed an increased sphericity and a loss in ellipticity in Itgb1-deficient MCs compared to wt controls, describing more spherical MC shape in the absence of Itgb1 (Fig. 1H, I). Additionally, the lack of Itgb1 in MCs also fundamentally affected the MC distribution in the mouse ear skin. In particular, the overall mean distance of MCs to the closest blood vessel was significantly increased in MC^ΔItgb1^ (Fig. 1J). The aberrant MC positioning was most striking in the blood vessel associated MC fraction with physical contact with the blood endothelium (Fig. 1K). In contrast to a homogeneous MC distribution in the mouse ear skin of MC^wt^ with numerous MCs aligning arterioles in high density, the perivascular MC positioning was fundamentally impaired. In fact, periarteriolar MC numbers were low or not detectable in MC^ΔItgb1^ mice. Interestingly, this effect was evident but less prominent along venules. (Fig. 1L, M). Quantification confirmed that the distance of MC^ΔItgb1^ to both arterioles and venules was increased, but with a much greater discrepancy at arterioles (Fig. 1N). In line, the density of periarteriolar MCs was decreased in MC^ΔItgb1^ compared to MC^wt^ (Fig. 1O). In contrast, ellipticity was decreased for both periarteriolar and perivenular MCs (Fig. 1P). Consistent with the publication by Kaltenbach et al. (16), MC distance to nearest neighbor was decreased in MC^ΔItgb1^ independent of the MC localization, indicating a cluster formation of mature skin MCs distant from blood vessels (Fig. 1Q). Collectively, Itgb1 is crucial for spindle-like morphology, perivascular alignment and correct dermal blood vessel positioning of skin MCs, especially at arterioles.

**Figure 1.**
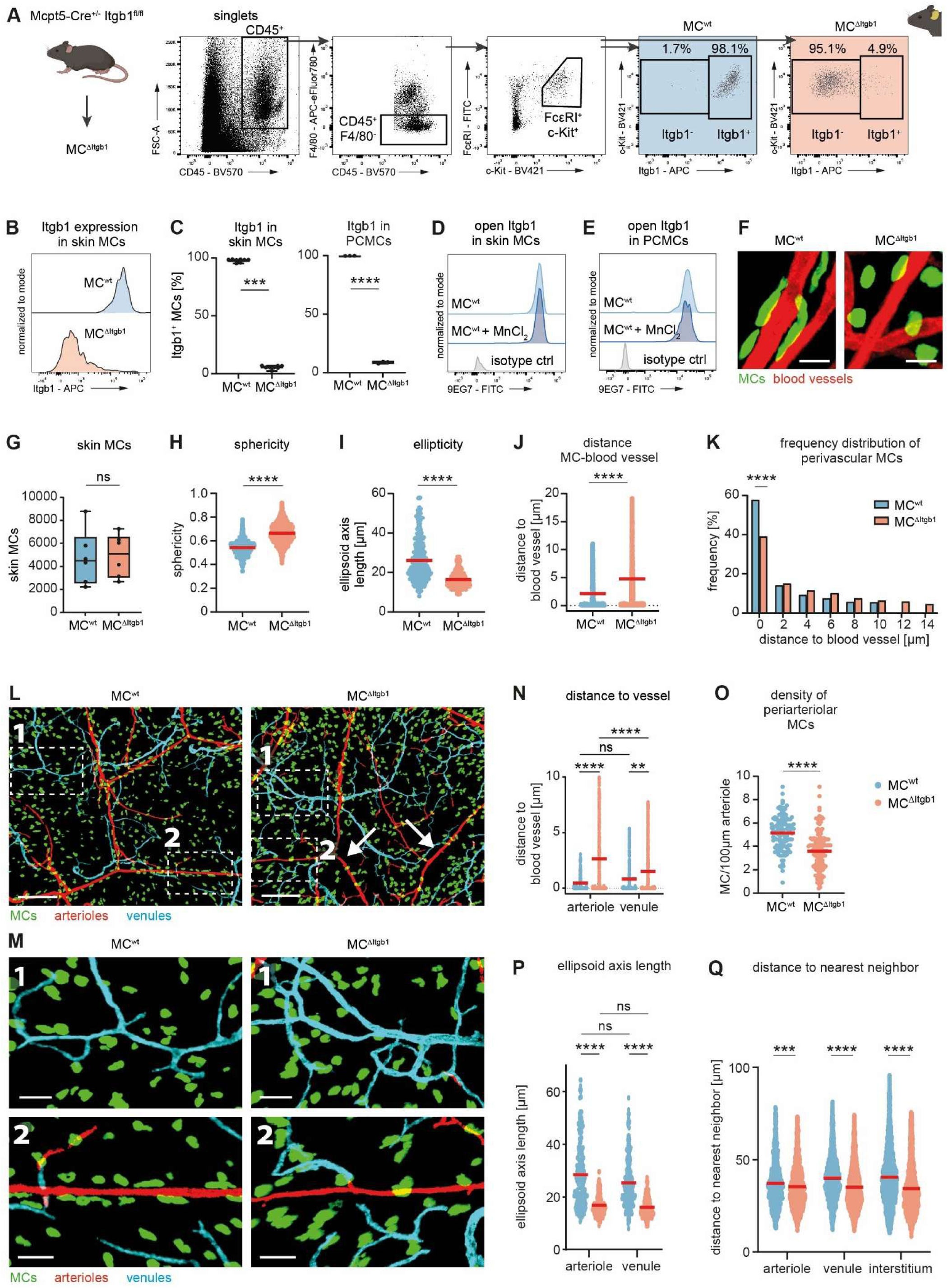
Elongated morphology, tissue distribution and blood vessel alignment of perivascular MCs depends on Itgb1. (A) To study the effect of Itgb1 in connective-tissue type MCs, Mcpt5-Cre^+/-^ Itgb1^fl/fl^ mice were used and skin MCs were gated in flow cytometry analysis as CD45^+^ F4/80^-^ FcεRI^+^ c-kit^+^ cells, based on FMOs. (B) Expression of Itgb1 as median fluorescence intensity (MFI) in skin MC^wt^ and MC^ΔItgb1^. (C) Quantification of Itgb1^+^ MCs in ear skin (n=8) and PCMCs (n=3). Active Itgb1 conformation in MC^wt^ was confirmed in (D) ear skin and (E) PCMCs by 9EG7 binding to the open formation. Negative control: isotype control; positive control: administration of 1mM MnCl_2_. (F) Representative confocal laser scanning microscopy images of MC^wt^ and MC^ΔItgb1^ ear skin samples for characterization of MC morphology. Visualization as maximum intensity projection (MIP). Red – tomato lectin DyLight 488, green – avidin Texas Red, scale bar 20µm. (G) MC counts in ear skin samples (n=6). Quantitative image analyses of (H) sphericity and (I) ellipticity as MC morphology parameters and (J) average distance and (K) frequency distribution of MCs to closest blood vessels (data points represent individual cells from N=3 mice per group with H: n=3017 (MC^wt^) and n=4639 (MC^ΔItgb1^), I: n=524 and n=547, J-K: n=2622 and n=4206). (L) Representative confocal laser scanning microscopy images of MC^wt^ and MC^ΔItgb1^ ear skin samples for quantification of MC-blood vessel characteristics. Overview images with (M) zoom-ins for (1) perivenular and interstitial, and (2) periarteriolar MCs. Arrows indicate low periarteriolar MC density in MC^ΔItgb1^ mice. Visualization as MIP. Red – tomato lectin DyLight 488, green – avidin Texas Red, turquois – endomucin (AF647). Scale bar 200µm (overview) or 50µm (zoom-in). Quantitative image analyses of (N) distance of MCs to arterioles or venules, (O) periarteriolar MC density, (P) ellipsoid axis length and (Q) distance to nearest neighbor as readout of MC density (data points represent individual cells from N=3 mice per group with N: n=143 to 274 (MC^wt^) and n=251 to 364 (MC^ΔItgb1^), O: n=130 and n=166, P: n=201 to 392 and n=187 to 358, Q: n=1028 to 3370 and n=1781 to 3887). Red bars indicate mean; ** p<0.01, *** p<0.001, **** p<0.0001, ns – not significant.

### Lack of integrin β1 in MCs results in drastically reduced skin inflammation upon CHS

We and others previously reported (9,20,21) that MCs are essential in the onset and kinetics of skin inflammation, e.g. in the mouse model of contact hypersensitivity (CHS). Recapitulating a fundamentally disturbed MC morphology and perivascular positioning in MC^ΔItgb1^ already in physiological conditions, we studied the influence on MC functions in the context of inflammation. In CHS, immune cell infiltration can be assessed as ear swelling upon epicutaneous application of DNFB in sensitized mice. While ear swelling peaked at 48h after challenge in MC^wt^ mice, almost no ear swelling was detected in MC^ΔItgb1^ mice, comparable to vehicle treated mice (Fig. 2A). Flow cytometry analysis at this time point revealed a significantly reduced number of leukocytes in the ear skin (Fig. 2B). In fact, cell counts of almost all analyzed leukocyte populations were reduced in the ear skin of MC^ΔItgb1^ mice 48h after DNFB, relative to MC^wt^ (Fig. 2C). In detail, while the frequency of each population from total infiltrated leukocytes remained unaffected (Fig. 2D), cell counts of neutrophils, monocytes, macrophages, dendritic cells (DCs) and CD8^+^ T cells were markedly reduced in absence of Itgb1 (Fig. 2E). Remarkably, at 12h after challenge, increased numbers of neutrophils, monocytes and DCs were found in the blood of MC^ΔItgb1^ mice (Fig. 2F-G) and an increased frequency of CD8^+^ T cells in the inguinal lymph node (LN) (Fig. 2H). Collectively, MC^ΔItgb1^ mice show an abolished ear swelling in CHS with decreased counts of leukocytes in the ear skin and increased counts in the blood circulation. These data suggest that the lack of Itgb1 in MCs affects the blood leukocytes’ capacity to extravasate to the skin tissue, but not their mobilization.

**Figure 2.**
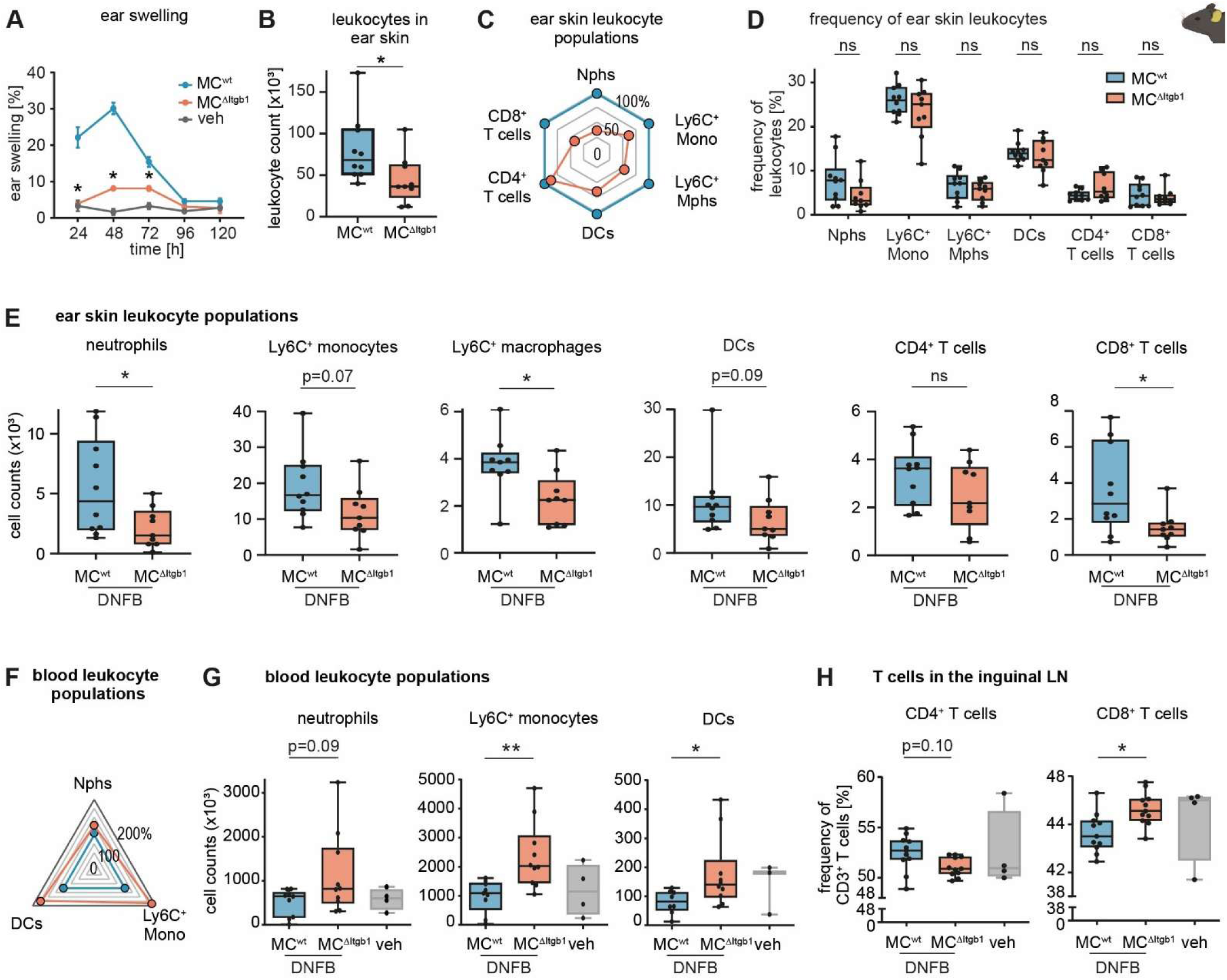
Absence of Itgb1 in MCs results in reduced ear swelling and ear skin inflammation upon CHS. The mouse model of contact hypersensitivity (CHS) evaluates the skin inflammation of sensitized mice upon the second encounter with the organic hapten 1-fluoro-2,4-dinitrobenzene (DNFB; challenge). (A) Ear swelling measurement as assessment of inflammation in MC^wt^ and MC^ΔItgb1^ mice upon challenge in CHS. Vehicle (veh) controls: solvent only-treatment. Measurements at indicated timepoints were normalized to values before treatment. Asterisks indicate significantly different results of MC^ΔItgb1^ to MC^wt^ (n=4 (DNFB); veh=3). (B) Flow cytometry analysis of total leukocyte counts in ear skin tissue 48h after challenge. (C) Relative counts of ear skin leukocyte populations at 48h after challenge. Values of MC^ΔItgb1^ were normalized to MC^wt^. (D) Frequency and (E) total counts of ear skin leukocyte populations at 48h after challenge (B-E: n=10 (MC^wt^) and n=9 (MC^ΔItgb1^)). (F) Relative counts of blood leukocyte populations at 12h after challenge. Values of MC^ΔItgb1^ were normalized to MC^wt^. (G) Total counts of blood leukocyte populations and (H) frequencies of T cells in inguinal lymph node (LN) at 12h after challenge (F-H: n=9 to 11 (MC^wt^), n=10 to 11 (MC^ΔItgb1^), and n=4 (veh)). * p<0.05, ** p<0.01 or p-values as indicated, ns – not significant. DCs – dendritic cells, Mono – monocytes, Mphs – macrophages, Nphs – neutrophils.

### Intravascular degranulation of perivascular MCs is drastically decreased in absence of integrin β1 *in vivo*

The abnormal vessel alignment and morphology of perivascular skin MCs, abrogated ear swelling and impaired leukocyte skin infiltration in MC^ΔItgb1^ mice might point to a defect in MC functionality. Remarkably, isolated skin MCs featured a reduced granularity in steady state in the absence of Itgb1, while the cell size of Itgb1-deficient MCs appeared increased (Fig. 3A-B). In the CHS model, IL-33 and ATP, two alarmins released by stressed keratinocytes, are critical for MC activation and degranulation (22). Of note, untreated MC^ΔItgb1^ mice showed both a reduced number of MCs expressing the IL-33 receptor ST2 and reduced expression level of ST2 on ST2^+^ MCs (Fig. 3C-D). Further, MC^ΔItgb1^ mice had reduced frequencies of MCs expressing the ATP receptor P2X7, while the overall expression level was unchanged (Fig. 3E-F). Interestingly, also the expression level of the stem cell factor (SCF) receptor c-kit was markedly reduced in MC^ΔItgb1^ (Fig. 3G). The collective results indicate a substantial impairment of receptor expression on skin MC^ΔItgb1^ (Fig. 3H). It is thus plausible that Itgb1-deficient MCs may exhibit impaired activation mechanisms. As previously described, perivascular MCs are capable of directional degranulation into the blood vessels via intraluminal cell extensions (14). We therefore studied the intravascular MC degranulation capacity *in vivo* upon epicutaneous DNFB treatment. Briefly, avidin binds to heparin in MC granules, thereby labeling exocytosed MC granules. Following the membrane disintegration during MC degranulation, intracellular granules will also be stained by avidin (14,23). Due to the i.v. application of fluorochrome-conjugated avidin upon DNFB application, particularly perivascular MCs with access to the blood stream can be labeled and detected by intravital 2-photon microscopy (Fig. 3I). Interestingly, both MC^wt^ and MC^ΔItgb1^ were capable of incorporating i.v. applied avidin (Fig. 3J). Selected close-up views revealed individual perivascular MCs forming intravascular sheets, irrespective of Itgb1 expression. However, the proportion of MC membrane surface inside the blood vessel was markedly reduced in MC^ΔItgb1^, compared to wt controls (Fig. 3K). In addition, the extent of avidin uptake was further analyzed by flow cytometry analysis of digested *Mcpt5-cre^+^ tdTomato^+^* MC reporter mouse ear skin (Fig. 3L). The number of avidin^+^ MCs was similar between MC^wt^ and MC^ΔItgb1^ (Fig. 3M). However, Itgb1-deficient MCs presented a marked reduction in median fluorescence intensity (MFI) for i.v. applied avidin compared to wt controls (Fig. 3N-O), suggesting an impaired intravascular degranulation efficiency in MC^ΔItgb1^ mice *in vivo*.

**Figure 3.**
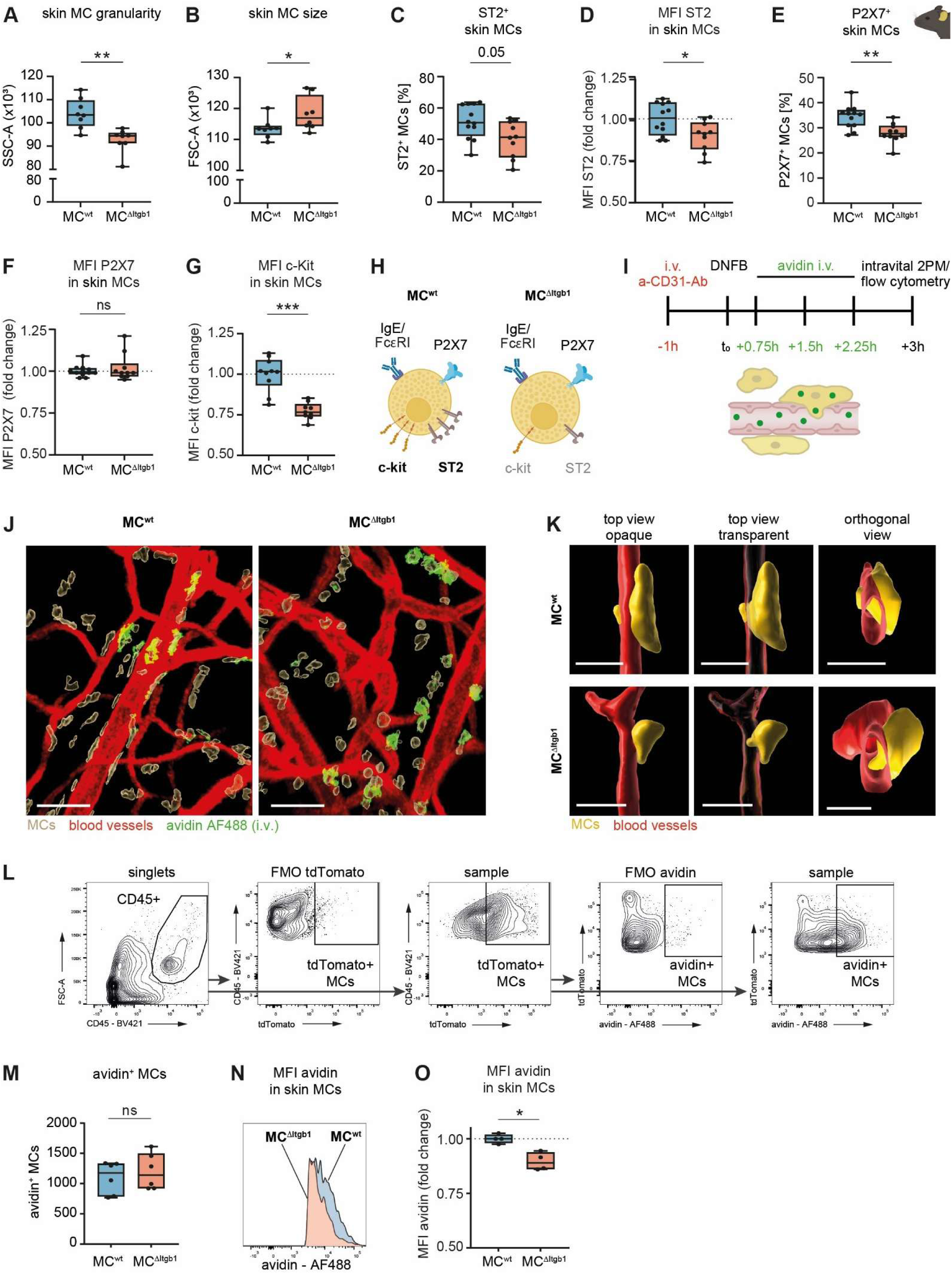
Altered MC granularity, receptor expression and decreased intravascular MC degranulation in the absence of Itgb1 *in vivo*. (A) Granularity and (B) size of ear skin MCs in MC^wt^ and MC^ΔItgb1^ mice (A-B: n=8). (C-D) Frequency of ear skin MCs and expression levels of IL-33 receptor ST2 and (E-F) ATP receptor P2X7 as median fluorescence intensity (MFI) (C-F: n=12 (MC^wt^) and n=10 (MC^ΔItgb1^). (G) Expression of c-kit as MFI on ear skin MCs. (H) Schematic overview of the expression levels of FcεRI, P2X7, ST2 and c-kit on skin MC^wt^ and MC^ΔItgb1^. (I) Experimental scheme for the quantification of intraluminal MC degranulation for analysis by intravital 2-photon microscopy (2PM) or flow cytometry. At least 3 types of MCs can be found in 2PM: non-vessel associated interstitial MCs, perivascular MCs without and with contact to the blood stream, with the latter capable of uptake of i.v. applied avidin. (J) Representative intravital 2PM images of MC^wt^ and MC^ΔItgb1^ ear skin samples depicting uptake of i.v. applied avidin by perivascular MCs. Visualization as maximum intensity projection (MIP) or 3D surface rendered objects. Red – CD31-AF647, beige – tdTomato, green – i.v. applied avidin AF488. Scale bar 50µm. (K) Representative intravital 2PM images of individual MCs with interaction with the blood stream in MC^wt^ and MC^ΔItgb1^ ear skin samples. Depiction of 3D surface rendered objects with opaque or transparent blood vessel structures (top view), and orthogonal view of the same cell, respectively. Red – CD31-AF647, yellow – tdTomato. Scale bar 20µm. (L) Gating strategy for detection of uptake of i.v. applied avidin by tdTomato^+^ MCs. MCs were gated as CD45^+^ tdTomato^+^ cells, based on FMOs. (M) Total counts of avidin^+^ MCs (n=6). Quantification of avidin uptake by MCs as (N-O) measured fluorescence intensity (n=4). MFI of MC^ΔItgb1^ was normalized to MC^wt^. * p<0.05, ** p<0.01, *** p<0.001, or p-values as indicated, ns – not significant.

### integrin β1 is essential for MC degranulation kinetics *in vitro*

In order to clarify whether the reduced avidin binding resulted from impaired directional intravascular degranulation or from generally impaired degranulation proficiency, we studied MC functionality in more detail by using PCMCs *in vitro*. Retrieved via non-enzymatic isolation, PCMCs represent a mature mast cell population that retains morphological, phenotypic and functional characteristics. They are therefore considered a suitable model for mature connective-tissue mast cells while preserving enzyme-sensitive surface receptors such as integrins and activation markers (24). To this end, non-adhesive PCMCs from MC^wt^ and MC^ΔItgb1^ mice were cultured in suspension without interaction with other cells or extracellular matrix, which allowed us to study intrinsic effects upon MC activation. Here, PCMCs were stimulated with ATP and IL-33 simultaneously, the two key alarmins activating MCs in the CHS model (22). Unspecific MC stimulation with the calcium ionophore A23187, or specific activation of anti-DNP-IgE sensitized PCMCs by FcεRI crosslinking with DNP-BSA served as controls. MC degranulation was assessed by means of CD107a expression using flow cytometry (for total numbers of degranulated MC) or by β-hexosaminidase release assay (to quantify MC degranulation capacity). While IL-33 alone did not induce degranulation *in vitro* (Fig. S2A), A23187, ATP, ATP+IL-33 and DNP/IgE stimulation resulted in a moderate degranulation which was unaffected by the absence of Itgb1, when degranulated PCMCs were analyzed as CD107a^+^ (Fig. 4A). Remarkably, when MC degranulation was assessed by β-hexosaminidase release assay, degranulation was reduced in Itgb1-deficient PCMCs compared to wt controls (Fig. 4B). In line with our *in vivo* findings, this data suggest that MCs are able to degranulate in absence of Itgb1, but show an aberrant extent of degranulation or granule release mechanism. Of note, unstimulated PCMCs displayed an increased expression level of P2X7 in the absence of Itgb1 (Fig. 4C), which was in contrast to a decreased expression in ear skin MCs (Fig. 3E). Expression levels of ST2 and frequency of P2X7^+^ and ST2^+^ MCs were similar between MC^wt^ and MC^ΔItgb1^ (Fig. 4D, Fig. S2B-C). Interestingly, unstimulated Itgb1-deficient PCMCs had decreased expression levels of FcεRI and c-kit (Fig. 4E-F). These findings led us hypothesize that Itgb1 may have an intrinsic effect on MC activation, probably further potentiated by the altered expression levels of FcεRI and P2X7 (Fig. 4G). Therefore, we examined *in vitro* PCMC degranulation in a high-resolution confocal laser scanning microscopy live-cell imaging approach (25,26). Briefly, untreated or anti-DNP IgE sensitized PCMCs were stimulated with ATP or DNP-BSA, respectively. The addition of fluorochrome-conjugated avidin to the supernatant resulted in binding to MC granules as soon as they were exocytosed, thereby allowing to track the course and kinetic of degranulation (Fig. 4H). We observed a rapid avidin accumulation on the surface of degranulating PCMCs, visualizing the granule budding during exocytosis. In order to quantify the process of degranulation, we decided to decrease the stimulus to 5µM ATP or 1ng/ml DNP-BSA, respectively. Time-lapse imaging visualized a rapid and strong avidin accumulation in MC^wt^ upon ATP stimulation, while avidin labeling was much less intense in MC^ΔItgb1^ (Fig. 4I, Video S1). Quantification of the avidin intensity of numerous individual PCMCs confirmed that MC^wt^ progressively accumulated avidin over 30min, while the avidin intensity in MC^ΔItgb1^ was significantly reduced (Fig. 4J). DNP/IgE stimulation also resulted in a moderate avidin accumulation in both MC^wt^ and MC^ΔItgb1^, with a slightly decreased avidin intensity in the later (Fig. 4K-L, Video S2). No differences in the cell size or granularity between MC^wt^ and MC^ΔItgb1^ were observed prior or post stimulation (data not shown). Of note, contents of released pro– and anti-inflammatory mediators, such as TNF, IFN, IL-5, IL-6 or IL-10, respectively, were similar in MC^wt^ and MC^ΔItgb1^ 6h after stimulation *in vitro* (Fig. S2D). Thus, PCMCs exhibited delayed degranulation kinetics, but not cytokine de novo synthesis, upon stimulation with ATP and DNP/IgE in the absence of Itgb1 *in vitro*.

**Figure 4.**
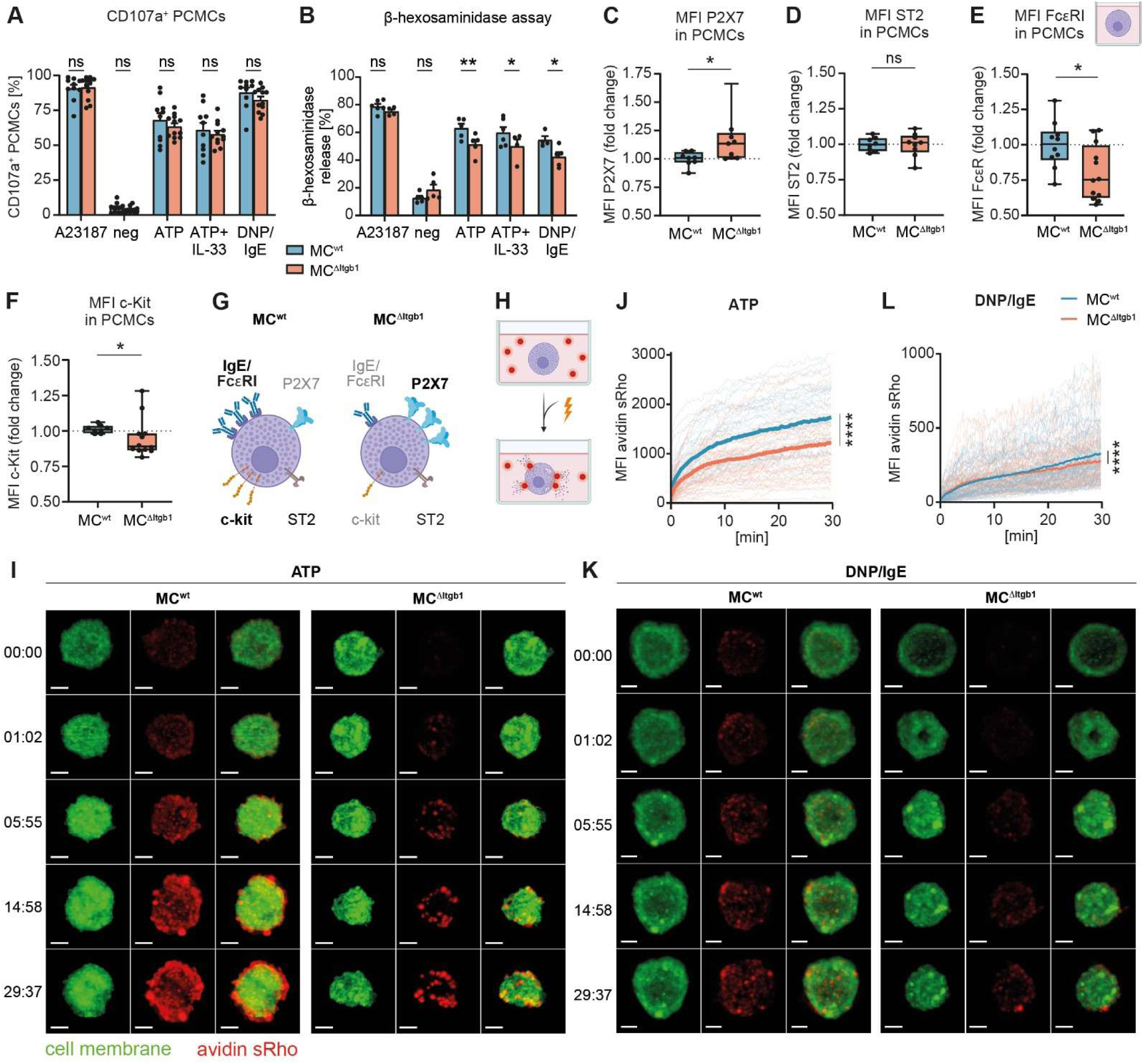
Impaired PCMC degranulation kinetics and receptor expression in the absence of Itgb1 *in vitro*. Quantification of PCMC degranulation upon stimulation with calcium ionophore A23187 (positive control), medium (negative control), 500µM ATP, 500µM ATP + 50ng/ml IL-33 or 100ng/ml DNP-BSA (sensitized with anti-DNP IgE), respectively, by (A) CD107a^+^ MCs in flow cytometry (n=10 (MC^wt^) and n=12 (MC^ΔItgb1^) or (B) β-hexosaminidase release assay (n=4-6 and n=4-5). Expression of (C) P2X7, (D) ST2, (E) FcεRI, (F) c-kit as median fluorescence intensity (MFI, C-D: n=8 (MC^wt^) and n=9 (MC^ΔItgb1^), E: n=10 and n=12, F: n=8 and n=12). MFI of MC^ΔItgb1^ was normalized to MC^wt^. (G) Schematic overview of the expression levels of FcεRI, P2X7, ST2 and c-kit on PCMCs from MC^wt^ and MC^ΔItgb1^ mice. (H) Experimental setup for live-cell stimulation and visualization of PCMC degranulation. (I) Representative images of *in vitro* live-cell confocal laserscanning microscopy of MC^wt^ and MC^ΔItgb1^ upon stimulation with 5µM ATP for 30min. Green – CellBrite steady 405, red – avidin sulforhodamine 101, visualization as maximum intensity projection (MIP) of single channels (left and middle) and merge (right). Scale bar 5µm. (J) Single cell (faint lines) and mean (bold line) avidin sulforhodamine 101 intensity of MC^wt^ and MC^ΔItgb1^ upon stimulation with 5µM ATP for 30min (lines represent individual cells from N=3 mice per group with n=36 (MC^wt^) and n=46 (MC^ΔItgb1^). (K) Representative images of *in vitro* live-cell confocal laser scanning microscopy of MC^wt^ and MC^ΔItgb1^ upon stimulation with 1ng/ml DNP-BSA of anti-DNP IgE sensitized PCMCs for 30min. Green – CellBrite steady 405, red – avidin sulforhodamine 101, visualization as MIP of single channels (left and middle) and merge (right). Scale bar 5µm. (L) Single cell (faint lines) and mean (bold line) avidin sulforhodamine 101 intensity of MC^wt^ and MC^ΔItgb1^ upon stimulation with 1ng/ml DNP-BSA of anti-DNP IgE sensitized PCMCs for 30min (lines represent individual cells from N=3 mice per group with n=74 (MC^wt^) and n=71 (MC^ΔItgb1^). * p<0.05, ** p<0.01, **** p<0.0001, ns – not significant. See also Video S1 and S2.

### integrin β1 is involved in intracellular signaling upon ATP stimulation

Given the impact of Itgb1 on MC degranulation kinetics, we next investigated the underlying intracellular signal transduction downstream of ATP stimulation (22). The PI3K-AKT pathway has been shown to mediate MC degranulation (28), whereas the SHIP1 pathway has been identified as a limiting regulator of MC degranulation (29). It is important to note that a balanced activity of the PI3K-AKT pathway and the SHIP1 pathway is crucial for determining the activation or inhibition of degranulation, respectively (Fig. 5A). While most studies focused on IgE stimulation in bone marrow-derived MCs (BMMCs), there is limited but suggestive evidence that SHIP1/PI3K-AKT signaling might also be involved upon ATP stimulation (30). By western blot analysis, we investigated the activation of SHIP1 and AKT, the latter of which is activated downstream of PI3K. While only weak phosphorylation of SHIP1 was detected in MC^wt^ upon ATP stimulation, a strong induction of pSHIP1 was observed in MC^ΔItgb1^. In contrast, both MC^wt^ and MC^ΔItgb1^ showed moderate phosphorylation of AKT (Fig. 5B). Indeed, pAKT peaked early in MC^wt^, with even stronger signals in MC^ΔItgb1^ (Fig. 5C). Further, it is noteworthy that strong phosphorylation of SHIP1 was detected in MC^ΔItgb1^ upon ATP stimulation, compared to low phosphorylation in MC^wt^ (Fig. 5D). Importantly, pAKT-to-pSHIP1 ratio was skewed in Itgb1-deficient PCMCs, demonstrating an imbalance towards the SHIP1 pathway (Fig. 5E). Specifically, normalization of MC^ΔItgb1^ to MC^wt^ uncovered a drastic shift towards pSHIP1 over pAKT in the absence of Itgb1 (Fig. 5F). These findings indicate that the PI3K-AKT pathway might be outcompeted by the strong SHIP1 pathway signaling. Next, we evaluated whether calcium (Ca^2+^) signaling might be causative for the strongly imbalanced SHIP1/PI3K signaling in MC^ΔItgb1^. Since P2X7 as an ATP-regulated ion channel mediates the Ca^2+^ influx upon activation (31), extracellular Ca^2+^ influx upon ATP stimulation was analyzed *in vitro*. Remarkably, extracellular Ca^2+^ influx was increased in MC^ΔItgb1^ compared to MC^wt^ (Fig. 5G). Quantification of peak Ca^2+^ influx and area under the curve confirmed an increased Ca^2+^ influx in Itgb1-deficient PCMCs (Fig. 5H-I). These findings are in line with an elevated P2X7 expression in MC^ΔItgb1^ (Fig. 4C). Of note, baseline Ca^2+^ influx was not affected (Fig. 5J). Previously, ATP was shown to induce the release of Ca^2+^ from intracellular stores (32) and store-operated Ca^2+^ entry (SOCE) was reported to be a dominant Ca^2+^ influx pathway in MCs during allergic airway inflammation induced by house dust mites (33). However, intracellular Ca^2+^ store depletion and subsequent SOCE were not dependent on Itgb1 (Fig. 5K-O). To assess the promoting effect of PI3K on MC degranulation upon ATP stimulation, a selective PI3K inhibitor was applied to PCMCs and degranulated MCs were quantified by flow cytometry. LY294002 at 50µM did not affect cell viability (data not shown), but resulted in a diminished degranulation upon stimulation with DNP/IgE or ATP in both MC^wt^ and MC^ΔItgb1^ (Fig. S2E). However, the inhibitory effect on MC degranulation upon ATP stimulation was more prevalent in MC^ΔItgb1^ compared to MC^wt^ (Fig. 5P-Q), which might be due to a predominant pSHIP1 inhibition in MC^ΔItgb1^. Thus, increased Ca^2+^ influx due to elevated P2X7 levels might contribute to insufficient signaling upon ATP stimulation in MC^ΔItgb1^ with a clear predominant effect of pSHIP1 over pAKT, leading to a stronger inhibition of degranulation *in vitro* (Fig. 5R).

**Figure 5.**
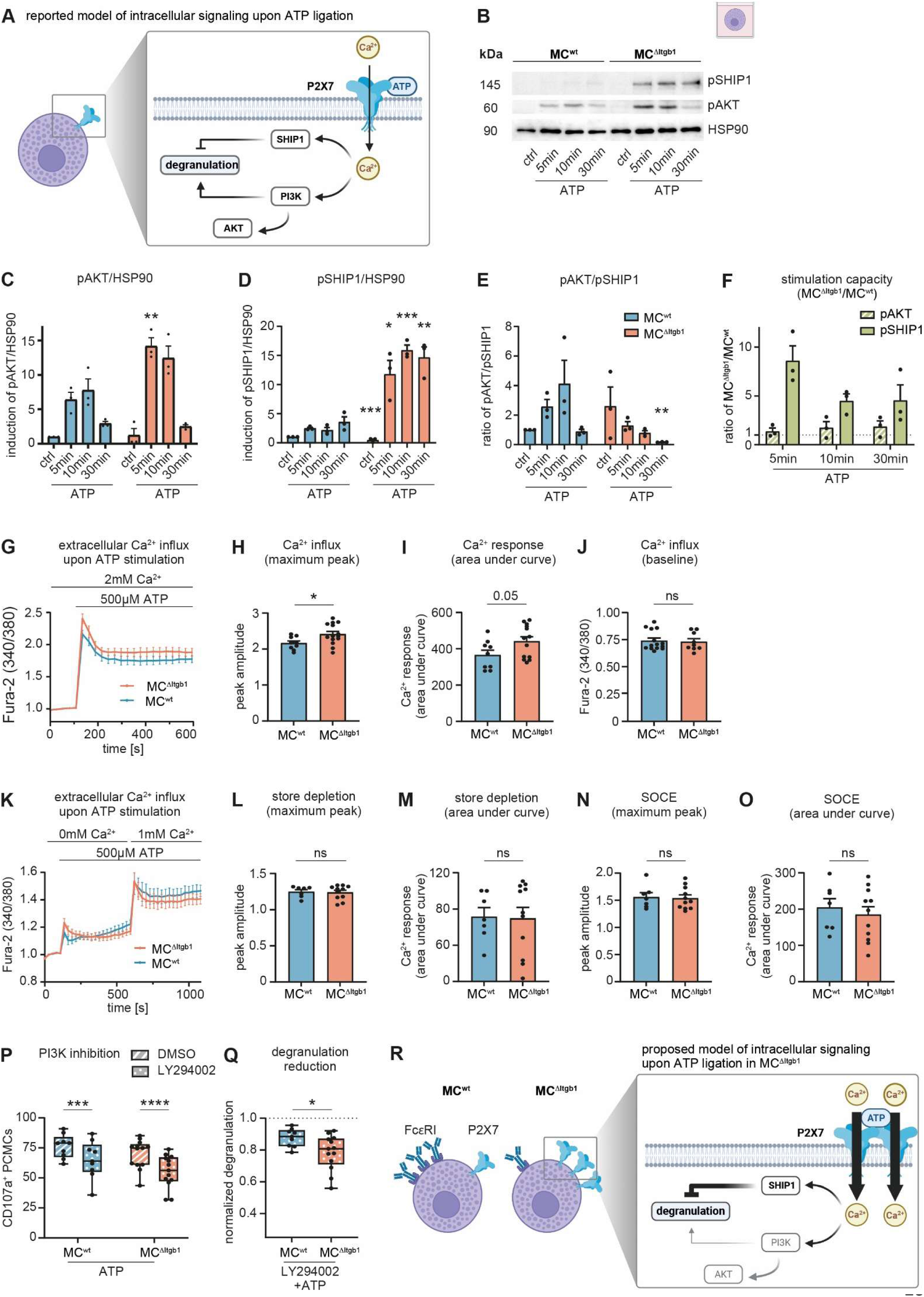
Abnormal intracellular phosphorylation patterns and calcium influx upon ATP stimulation in Itgb1-deficient PCMCs *in vitro*. (A) Schematic description of reported model for intracellular mechanisms affecting degranulation in MCs. Calcium (Ca^2+^) influx upon P2X7 ligation by ATP initiates phosphorylation of SHIP1 and AKT (activated downstream of PI3K), exhibiting either an inhibiting or activating effect on MC degranulation, respectively (28,29). (B) Western blot analysis of phosphorylated (p) SHIP1 and AKT in unstimulated (ctrl) MC^wt^ and MC^ΔItgb1^ or upon stimulation with 500µM ATP for up to 30min. Loading control: heat shock protein (HSP) 90. Ratio of (C) pAKT to HSP90, (D) pSHIP1 to HSP90 and (E) pAKT to pSHIP1, with normalization of all values to MC^wt^ ctrl. (F) Ratio of pAKT and pSHIP1 in MC^ΔItgb1^ normalized to MC^wt^, (C-F: based on (B), data points represent individual experiments with n=3). (G) Ca^2+^ influx as Fura-2 emission (at 340/380nm) of MC^wt^ and MC^ΔItgb1^ in 2mM Ca^2+^ for 10min. Baseline measurement for 108s, before stimulation with 500µM ATP was initiated. Normalization of values to mean of baseline of MC^wt^ and MC^ΔItgb1^, respectively. Quantification of (H) peak Ca^2+^ influx, (I) area under curve and (J) non-normalized Fura-2 baseline signal, based on (G) (G-J: n=9 (MC^wt^) and n=13 (MC^ΔItgb1^)). (K) Ca^2+^ influx as Fura-2 emission (at 340/380nm) of MC^wt^ and MC^ΔItgb1^ in 0mM Ca^2+^ for 20min. Baseline measurement for 108s, before stimulation with 500µM ATP (store depletion), followed by the supply of 2mM Ca^2+^ (resulting in 1mM Ca^2+^ in the medium) after 621s (store-operated Ca^2+^ entry, SOCE). Normalization of values to mean of baseline of MC^wt^ and MC^ΔItgb1^, respectively. Quantification of (L) peak Ca^2+^ influx and (M) area under curve upon store depletion, and (N) maximum peak and (O) area under curve upon Ca^2+^ application (SOCE) based on (K) (K-O: n=7 (MC^wt^) and n=11 (MC^ΔItgb1^)). (P) Inhibition of MC degranulation upon stimulation with 500µM ATP, after treatment with PI3K inhibitor LY294002 or DMSO for 30min. (Q) Normalization of MC degranulation to DMSO treatment upon stimulation with 500µM, based on (P) (P-Q: n=9 (MC^wt^) and n=13 (MC^ΔItgb1^). (R) Schematic model for the expression of FcεRI and P2X7 on MC^wt^ and MC^ΔItgb1^ and proposed impaired Ca^2+^ influx and phosphorylation pattern in MC^ΔItgb1^, resulting in reduced degranulation. * p<0.05, *** p<0.001, **** p<0.0001, ns – not significant.

### Lack of integrin β1 in MCs results in diminished blood vessel activation and neutrophil extravasation

Considering the impaired MC degranulation efficiency in the absence of Itgb1 *in vitro* and *in vivo*, their affected blood vessel alignment and the abrogated inflammation upon DNFB *in vivo*, we assumed that the Itgb1-mediated positioning of MCs is prerequisite for their vasoactive and leukocyte recruiting functions. Since MCs are in close relation with blood vessels (Fig. 1F), we investigated the activation of blood endothelial cells (ECs) upon CHS (Fig. 6A). Indeed, 12h post DNFB, ECs presented an upregulation of intercellular activation molecule 1 (ICAM-1) in MC^wt^ mice upon DNFB challenge, as compared to vehicle treated mice, that was abrogated in MC^ΔItgb1^ mice (Fig. 6B). The levels of vascular cell adhesion molecule 1 (VCAM-1) and E-selectin, two other activation markers on ECs required for leukocyte extravasation (34), were not altered at that time point (Fig. 6C-D). In addition, the surface expression of CD11b on circulating neutrophils, which is a marker for neutrophil activation and intravascular priming, was found significantly reduced in MC^ΔItgb1^ mice compared to MC^wt^, to a level even below the vehicle controls (Fig. 6E-F). Importantly, CD11b surface expression was inversely related to the blood neutrophil counts (Fig. 6G). Since neutrophils are the first immune effector cells recruited to inflammatory sites and crucial for the recruitment of further leukocyte populations (35–37), we focused on the study of neutrophil extravasation behavior in MC^ΔItgb1^ mice upon DNFB. Applying intravital 2-photon microscopy allowed us to monitor neutrophil rolling, adhesion and extravasation (14). Interestingly, neutrophil rolling flux and velocity were increased in MC^ΔItgb1^ mice compared to MC^wt^ mice, but fewer neutrophils showed firm adhesion at the endothelial walls (Fig. 6H-K, Video S3). Of note, microvascular and hemodynamic parameters were not different between MC^wt^ and MC^ΔItgb1^ mice (Fig. S3A-D). In line, markedly fewer neutrophils extravasated from the blood into the ear tissue when MCs lacked Itgb1 (Fig. 6L, see also Fig. 2E). Thus, neutrophil extravasation behavior was impaired in MC^ΔItgb1^ mice, potentially due to an abrogated activation of both circulating neutrophils and ECs.

**Figure 6.**
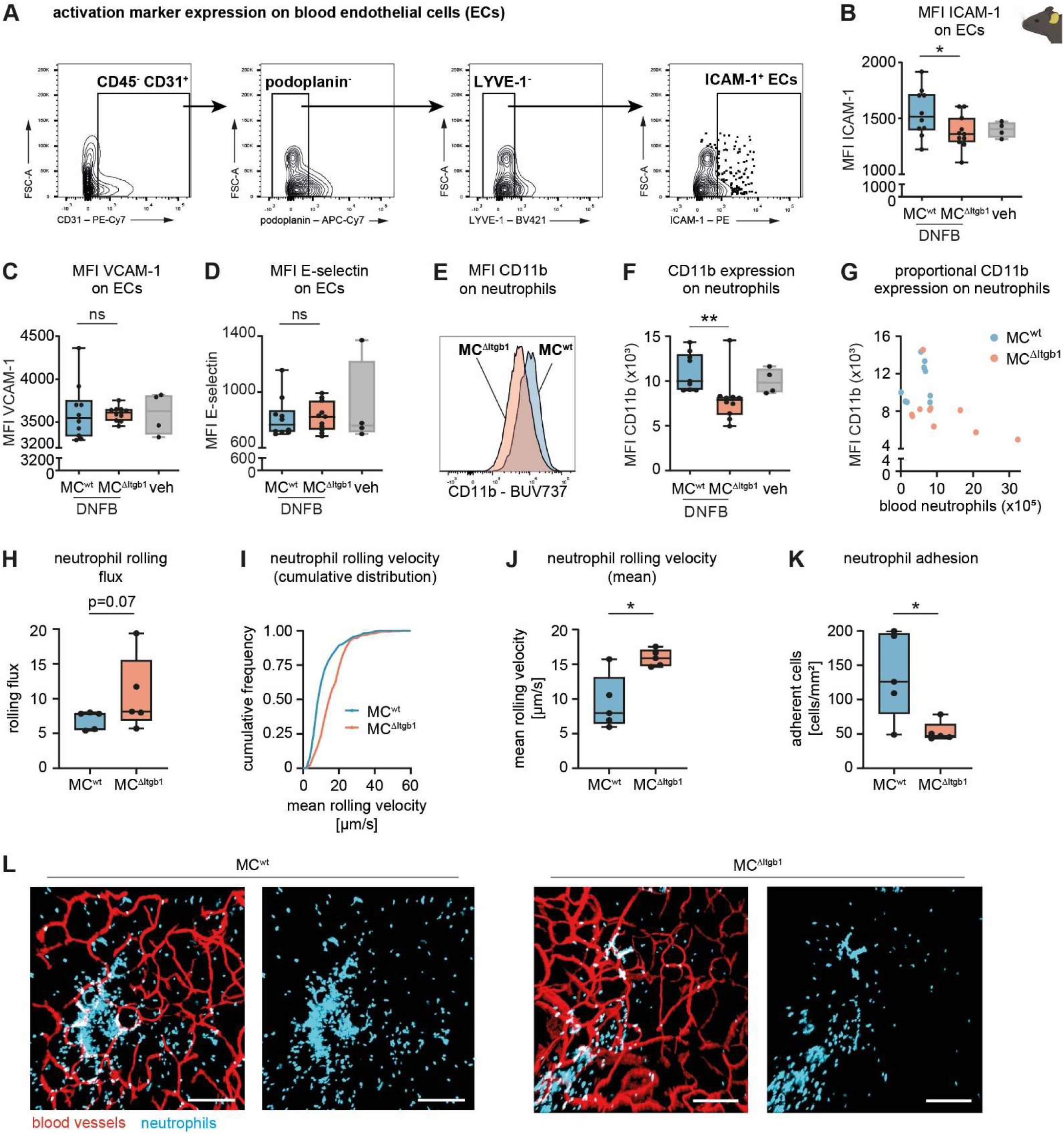
Reduced activation of blood endothelial cells and neutrophils results in attenuated neutrophil rolling and extravasation in MC^ΔItgb1^ mice. (A) Gating strategy for the identification of activation markers, such as intercellular adhesion molecule 1 (ICAM-1) on blood endothelial cells (ECs). ECs were gated as CD45^-^ CD31^+^ podoplanin^-^ LYVE-1^-^ cells, based on FMOs. Expression of (B) ICAM-1, (C) vascular cell adhesion protein 1 (VCAM-1) and (D) E-Selectin as median fluorescence intensity (MFI) on ECs in the ear skin and CD11b on blood neutrophils of MC^wt^ and MC^ΔItgb1^ mice 12h after DNFB treatment, respectively (E-F). Treatment with the solvent only served as vehicle (veh) controls (n=10 (MC^wt^) and n=11 (MC^ΔItgb1^), veh=4). (G) CD11b expression in relation to blood neutrophil counts, based on (F). (H) Mean number of rolling neutrophils, (I) cumulative and (J) mean neutrophil rolling velocity, and (K) mean number of adherent neutrophils in postcapillary venules of MC^wt^ and MC^ΔItgb1^ mice 12h after DNFB treatment (n=35 (MC^wt^) and n=37 (MC^ΔItgb1^) vessels of N=5 mice per group). (L) Representative intravital 2PM images of MC^wt^ and MC^ΔItgb1^ ear skin samples depicting neutrophil infiltration 12h after DNFB treatment. Visualization as maximum intensity projection (MIP) of single channel for Ly6G-PE (right) and merge (left). Red – CD31-AF647, cyan – Ly6G-PE. Scale bar 200µm. * p<0.05, ** p<0.01, or p-values as indicated, ns – not significant. See also Video S3.

Collectively, we discovered that the adhesion molecule Itgb1 is essential for MC localization and positioning in the skin perivascular niches, and for their alignment at the blood vessels. Further, Itgb1 is crucial for MC degranulation properties *in vivo*, thereby being indispensable for the MC-mediated vasoactive effects contributing to the activation of EC and neutrophil extravasation upon skin inflammation. We further identified impaired degranulation mechanisms in Itgb1-deficient MCs *in vitro*, suggesting an additional intrinsic role of Itgb1, which is essential for MC degranulation kinetics.

## DISCUSSION

MCs are strategically positioned at host interfaces that are exposed to environmental influences. Here, they are particularly closely associated with blood vessels, neurons, and hair follicles (11). As demonstrated recently, integrins, and in particular integrin β1 (Itgb1), are essential for MC positioning in the ear skin (16,18). Using the mouse line *Mcpt5-Cre Itgb1^fl/fl^*, we confirmed that Itgb1 is critical for the MCs’ tissue positioning in mouse ear skin, the alignment at blood vessels and the elongated morphology in the perivascular niche. Importantly, we further demonstrated a critical function of Itgb1 for MC activation and degranulation. Due to this dual function, Itgb1 is essential for MC-mediated activation of blood endothelial cells and initiation of leukocyte infiltration during skin inflammation.

The depletion of the β1 subunit abolishes the formation of various integrin heterodimers that are involved in binding to extracellular matrix (ECM) components (15,16). Binding to laminin (via α3β1, α6β1, α7β1), fibronectin (α4β1, α5β1) and collagen type IV (α1β1) is crucial for MC anchorage to the ECM and other cells, but may also be involved in MC survival and effector function, as reviewed in (15). Arterioles exhibit circumferential coverage from multiple layers of vascular smooth muscle cells (VSMCs). VSMCs, in turn, produce ECM molecules (38,39) that may mediate perivascular MC binding. Further, the arterioles’ basement membrane is rich in collagen (predominantly type I and IV) and laminin (40). Compared to arterioles, the ECM composition around venules is looser with only microdomains of laminin and collagen IV (41), accompanied by an incomplete coverage with VSMCs (42). In contrast, capillaries are marked by a complete absence of VSMCs (43). Importantly, Kaltenbach et al. recently described a close interaction of dermal MCs with arteriolar VSMCs and a type of periarteriolar fibroblasts, both expressing high amounts of the MC growth factor stem cell factor (SCF, also known as KIT ligand). Further, periarteriolar MCs displayed a marked upregulation of the MC protease Mcpt-6, even in the absence of integrins. Thus, the authors suggest that the SCF-rich environment in the perivascular niche may facilitate local MC maturation (16), which critically depends on SCF/c-kit signaling (44).

On the contrary, our results imply a defective MC morphology and phenotype, but also function, degranulation kinetics and intercellular communication in the absence of Itgb1. We unveiled that Itgb1 essentially affects the granularity and enhances the degranulation capacity, as demonstrated by reduced avidin uptake upon CHS in MC^ΔItgb1^ mice. Blood vessel-associated MCs form extensions into the vessel lumen, facilitating rapid acquisition of circulating IgE (14,45). In the absence of Itgb1, perivascular association and passive blood sampling is abrogated, as shown by Link et al (18). Itgb1 is critical for the formation of membrane protrusions in fibroblasts (46). In our study, however, we demonstrate that perivascular MCs, if positioned close enough to the blood vessels, are capable of penetrating the endothelial walls, independent of Itgb1. This finding suggests that other mechanisms might be involved in the MCs passage through the endothelial barrier that compensate the lack of Itgb1.

Recently, we visualized elongated blood vessel-associated MCs with intraluminal protrusions by which they directionally degranulate into the blood circulation (14). The here presented findings suggest that reduced intraluminal degranulation in Itgb1-deficient MCs may be due to several reasons: (1) a greater distance of perivascular MCs to blood vessels, (2) a smaller contact area of perivascular MCs due to reduced cell elongation, but also (3) impaired priming of perivascular MCs in the absence of Itgb1, or a combination of the mentioned options.

It has been published that integrin binding to ECM compartments in MCs transmits extracellular forces *in vivo* (47), Thus, reduced MC elongation and therefore mitigated interaction surface with adjacent cells (endothelial cells, VSMCs, or fibroblast) and ECM might essentially affect MC differentiation, which requires membrane-bound SCF on cells in the perivascular niche (48). Our finding of reduced c-kit expression on dermal MCs in the absence of Itgb1 *in vivo* supports the hypothesis of an impaired MC adhesion and priming. This is accompanied by lower expression of the ATP receptor P2X7 and the IL-33 receptor ST2, suggesting a link to a reduced MC activation upon inflammatory insult *in vivo*.

In order to rule out bystander effects of other cells which might affect MC functions, we investigated the role of Itgb1 in MC activation and degranulation *in vitro* using PCMCs which are growing non-adherent in suspension. Due to the mild isolation procedure of peritoneal lavage, native receptor expression is preserved and *ex vivo* activation is minimized. Thus, PCMCs are considered to closely reflect their physiological state (24). Even though cultured in the same medium containing equal amounts of SCF, MC^wt^ and MC^ΔItgb1^ exhibited distinct phenotypes and activation behavior. Thus, we demonstrated that Itgb1 has an intrinsic effect on the capabilities of MCs to secrete granules, independently of their attachment to adjacent cells or ECM. This let us hypothesize that Itgb1 interferes with intrinsic signaling pathways independent of adhesion-induced outside-in-signaling. Integrins physically interact with growth factors, in particular with receptor tyrosine kinases (RTKs), such as c-kit (49). Importantly, c-kit has been reported to impact on integrin binding in hematopoietic cells (50,51). However, how integrins affect c-kit functionality specifically in MCs is still not fully understood. It has been reported that c-kit and α4 integrin signaling is linked to the PI3K/AKT pathway in bone marrow-derived MCs (BMMCs) (52). Importantly, inhibition of the PI3K/AKT pathway resulted in deficient maturation of BMMCs and impaired expression of c-kit and FcεRI (53). In our study, we demonstrated reduced expression of c-kit and FcεRI of MC^ΔItgb1^ compared to MC^wt^ *in vitro*. This was coinciding with increased expression of the ATP receptor P2X7 in Itgb1-deficient PCMCs. MCs express different P2X receptors, with P2X7 as the main purinergic ion channel (54). P2X7 ligation by ATP initiates Ca^2+^ influx (31), leading to MC activation and degranulation (55,56). While the effect of Ca^2+^ influx on activation of PI3K – as a promotor of degranulation upon ATP or FcεRI-mediated stimulation by phosphorylation of Akt – is well studied in BMMCs (27,57), the effect in PCMCs is poorly understood. Here, we demonstrate for the first time that ATP-mediated stimulation induces rapid phosphorylation of SHIP1 and Akt in PCMCs. Remarkably, phosphorylation dynamics were abnormal in Itgb1-deficient PCMCs. More importantly, we demonstrated a profound imbalance in pAKT/pSHIP1 in MC^ΔItgb1^, which in turn might be associated with reduced degranulation. These findings suggest Itgb1 as a possible link for the expression and effector function of c-kit, FcεRI and P2X7, possibly via the PI3K/AKT pathway. Of note, direct interaction of Itgb1 with P2X7 or indirect association via membrane microdomains is currently unknown. Considering the fact that actin remodeling is linked to P2X7 receptor membrane trafficking and function (58), and the reported interactions of P2X7 and integrins β3 and β2 in other cells (59,60), it is thus plausible that there might also be a direct or indirect structural or functional association of Itgb1 and P2X7.

In line with the impaired MC receptor expression, we confirmed substantial defects in MC degranulation kinetics in the absence of Itgb1 in PCMCs by *in vitro* live cell imaging. Upon stimulation by ATP, but also DNP/IgE, MC degranulation was drastically impaired as demonstrated by reduced avidin binding in the context of MC granule release. These findings underline an essential functional link of Itgb1 to Ca^2+^ influx, affecting MC degranulation *in vitro*. More importantly, we confirmed impaired MC degranulation also *in vivo* upon CHS. Of note, prestimulation of BMMCs with IL-33 (57,61) or c-kit (62) was shown to potentiate the MC cytokine production upon ATP stimulation. Thus, reduced expression of the IL-33 receptor ST2 and c-kit in the absence of Itgb1 might contribute to reduced induction of inflammation mediated by perivascular MCs *in vivo*. Together, our findings demonstrate that in addition to its role in mediating the positioning, adhesion and elongation of perivascular MCs to ensure priming effects of SCF and activation by IL-33 and ATP, Itgb1 is also involved in intracellular signal transduction towards an effective Ca^2+^ signaling and degranulation. Thus, we reveal an intrinsic Itgb1 effect mediating dual functionality which is essential for both MC positioning and degranulation upon activation.

Importantly, absence of Itgb1 in MCs abrogated the ear swelling, but also skin infiltration of several leukocyte populations upon CHS. Together with the fact that PCMC cytokine release is not altered *in vitro*, this suggests a general impairment of MC degranulation or their specific capacity to activate the blood endothelium. Upon induction of inflammation, MC orchestrate vasodilatation and increased blood vessel permeability by release of histamine and TNF, eventually inducing leukocyte rolling and extravasation (14,36,63,64). Following inflammatory responses, neutrophils are rapidly recruited, initiating the onset of inflammation by subsequent infiltration of other leukocytes, such as monocytes and macrophages (36,37). The reduced upregulation of both CD11b on neutrophils, as surrogate for impaired neutrophil priming, and ICAM-1 on blood endothelial cells in MC^ΔItgb1^ mice upon CHS finally resulted in impaired neutrophil adhesion and extravasation. Thus, in the absence of Itgb1, MCs are positioned not close enough to the blood vessels to transmit the vasoactive effect efficiently priming the endothelial cells for full onset of inflammation. Consequently, our findings demonstrate that the dual role of Itgb1 in MC positioning and effective degranulation is ultimately indispensable for their communication with blood vessel endothelium, vasoactive functions and the initiation of leukocyte infiltration. This, in turn, is essential, for the efficient onset of the inflammatory response.

The observed Itgb1-dependent close alignment of MCs at arterioles might further facilitate systemic signal transduction via the blood circulation. In humans, severe hypersensitivity reactions, such as anaphylaxis, can be fatal, due to the rapid onset of systemic vasodilatation and the effect on many body systems. Up to date, epinephrine represents the major cornerstone of treatment for anaphylactic shock, as it is the primary agent shown to effectively restore and stabilize blood pressure (65). Despite symptomatic treatment and the off-label use of biologicals to treat anaphylaxis (66), there is a lack of proactive treatment of the causative initial effects of intraluminal MC degranulation. Overall, our findings underscore the central role of MCs in inflammatory responses due to their strategic location in the perivascular niche, from which they coordinate inflammatory responses and intercellular communication. By identifying the essential relevance of Itgb1 in this process, we are making a decisive advance in deciphering the molecular mechanism of perivascular MC adhesion and intravascular degranulation in order to discover potential targets for the treatment of excessive MC activation.

## MATERIALS AND METHODS

### Mice

*Mcpt5-Cre* mice (B6-Tg(Cma1-cre)#Roer (67)) were crossed to *b1-FL* mice (Itgb1^tm1Ref^ (19)) provided by Prof. R. Fässler, (Max-Planck-Institute, Munich). For intravital microscopy experiments, this mouse line was crossed to *tdTomato* mice (B6.Cg-Gt(ROSA)26Sor^tm9(CAG-tdTomato)Hze^/J (68)), resulting in a *Mcpt5-cre x tdTomato x b1-FL* mouse line (B6-Tg(Cma1-cre)#Roer x B6.Cg-Gt(ROSA)26Sor^tm9(CAG-tdTomato)Hze^/J x Itgb1^tm1Ref^). All mice had a C57BL/6 background and were housed in the Central Animal Facility, Otto von Guericke University Magdeburg, under specific pathogen-free conditions. All experiments were performed on mice aged 8-20 weeks, in accordance with institutional guidelines on animal welfare and were approved by the Landesverwaltungsamt Sachsen-Anhalt (42502-2-1732 UniMD). Litter mates were used as controls.

### Preparation and imaging of dermal sheet samples

For blood vessel staining, mice were anesthetized by i.p. injection of a mix of 1% ketamine (#1202, cp pharma) and 1% xylazine (#1320422; total 10µl/g BW) and fluorescently labeled anti-CD31 antibody or lycopersicon esculentum (tomato) lectin (VectorLabs) was injected i.v. After 5min circulation, mice were sacrificed and ears were isolated and split into dermal sheets. Unless stated differently, all following steps were performed at room temperature (RT) and mildly shaking with PBS as stock medium. After fixation in 1% paraformaldehyde (4°C, overnight) and permeabilization with methanol/acetone (1:1) for 10min each side on ice, samples were washed (PBS, 10min). For fluorescence signal enhancement, samples were incubated in 100mM glycine for 30min. After washing, blocking (0.5µg/ml streptavidin, 60min; #016-000-113, Jackson ImmunoResearch) and MCs staining with AF488-conjugated (#A21370, ThermoFisher) or TexasRed-conjugated (#A820, ThermoFisher) avidin (5µg/ml, 60min) was performed, followed by 4x washing. If antibody staining was required, blocking was performed by blocking buffer (2% serum matching the secondary antibody, 50mM glycine, 0.05% Tween 20, 0.1% Triton X-100, 1% bovine serum albumin, 60min). Subsequently, primary antibody staining took place in blocking buffer overnight (4°C), followed by 4x washing in 0.1% Tween 20, staining with a secondary antibody in 0.1% Tween 20 (60min, RT) and 4x washing. Samples were allowed to air dry before being mounted onto an object slide for microscopy in Vectashield Antifade Mounting Medium (#VEC-H-1000, Biozol), sealing the covering slip with nail polish. Imaging was performed within 1 to 4 days using an inverted confocal laser scanning microscope TCS SP8 (Leica Microsystems, Wetzlar, Germany). Illumination was performed via an HC PL APO 20×/0,75 IMM CORR CS2 oil dipping lens. If required, sequential scanning was applied to reduce bleed-through of fluorescence signal into adjacent channels. Images were recorded as 8-bit z-stacks with a resolution of 1024×1024 pixels with depth of 50-95µm and a z spacing of 3-5µm. For visualization and quantification, 8 to 12 images were acquired of randomly chosen regions of interest (ROI) of the whole sample. For tile scans, z-stacks with a resolution of 512×512 pixel were acquired and were stitched together, resulting in a viewing area from 1500-3500µm. For analysis, single images were visualized as surface rendered z-stacks. Raw data was analyzed and quantified using Imaris software (version 9.3, Oxford Instruments, Zurich, Switzerland).

### Contact hypersensitivity

For sensitization, mice were treated with 100µl 0.5% DNFB (v/v, #D1529, Sigma) in olive oil/acetone (1:4) on the shaved back skin. 6d later, mice were challenged with 0.2% DNFB on the ear, 10µl each side. Vehicle controls were treated with the solvent only. Ear thickness was measured before and at indicated time points after challenge with a micrometer (Mitutoyo, Takatsu-ku, Japan). For the assessment of skin inflammation, ear swelling was calculated as percent increase compared to pre-challenge ear thickness. For avidin uptake, see adapted CHS model below.

### Avidin uptake by perivascular MCs

Analysis of intraluminal MC degranulation was conducted as described before (14). Briefly, unsensitized mice were anaesthetized by i.p. injection of a mix of 1% ketamine (#1202, cp pharma) and 1% xylazine (#1320422; total 10µl/g BW). For intravital microscopy, blood vessels were stained by i.v. injection of anti-CD31 AF647 (#102416, BioLegend, 0.8µg/g BW) 20min prior to the CHS response which was induced by the application of 0.2% DNFB on the ear (10µl each side) on unsensitized mice. To inhibit vessel permeabilization, pyrilamine (#P5514, Sigma-Aldrich, 10µg/g BW), an H1 histamine receptor antagonist, was injected i.v. 20min before, and 1h and 2.5h after DNFB. AF488-conjugated avidin (#A21370, ThermoFisher, 2.4µg/g BW) was i.v. applied 3× every 45min after DNFB. Avidin accumulation in perivascular MCs was evaluated 3h post DNFB by intravital 2-photon microscopy (2PM) or flow cytometry (see below). Intravital 2PM was conducted as described before (14). Briefly, mice were anaesthetized by i.p. injection of ketamine/xylazine (10µl/g BW, see above). For continuous anesthesia, half-dose injection was repeated, if necessary. Intravital 2PM was performed with a Zeiss LSM 710 multiphoton laser scanning microscope with five non-descanned detectors. Excitation was performed with a MaiTai DeepSee (track 1) or an InSight X3 (track 2) laser (both by SpectraPhysics) at 840nm (8-14%) and at 980nm (2.2-2.4%), respectively. Illumination was realized via a x20 water-dipping lens with 1.0 NA. Second harmonic generation (SHG) signal (<485nm, short pass filter 485nm, track 1) was used to visualize skin tissue architecture. Anti-CD31 AF647 was detected with a bandpass (BP) filter (675/70nm, track 1), tdTomato-expressing MCs with a BP filter (617/73nm) and avidin AF488 with a BP filter (525/50nm, all track 2). Images were recorded as 12-bit z-stacks with a resolution of 512×512 pixel, covering a viewing area of 425×425μm in xy and a depth from 50-130μm with a z-spacing of 3-5μm.

### Analysis and quantification of neutrophil motility behavior

For the analysis of neutrophil motility behavior, unsensitized mice were treated with 0.2% DNFB on the ear (10µl each side). 8-12h post DNFB, intravital 2PM was performed with an upright fixed stage Leica Stellaris 8 DIVE multiphoton laser scanning microscope with 3 hybrid detectors and an 8kHz resonant scanner (Leica Microsystems, Wetzlar, Germany). For this, mice were anesthetized using isoflurane (1.5-2.0% in oxygen) via mask. Anti-CD31 AF647 (#102416, BioLegend, 1.5µg/g BW) and anti-Ly6G PE (#127608, BioLegend, 7.5µg) was applied by an i.v. co-injection. After technical setup and right before imaging, 50µl Fluoresbrite YG microspheres (1µm, #17154-10, PolySciences; 1:100 in PBS) for the evaluation of hemodynamic parameters were i.v. injected. Excitation was performed with a MaiTai DeepSee laser (SpectraPhysics) at 800nm (4.0-5.5%) via a x25 water-dipping lens with 1.0 NA. YG was detected at 440-500nm, PE at 535-575nm and AF647 at 650-700nm. 12-bit images were recorded with a resolution of 512×512 pixel. Neutrophil motility behavior was monitored using time lapse videos. Neutrophil rolling and adhesion were analyzed as previously described (69) on a single plane image (25fps for 1min) with a viewing area of 150×150µm at 8-10h post DNFB. Neutrophil behavior and hemodynamic parameters were analyzed manually over 1min, using FIJI software (70). Static images were analyzed and assessed using Imaris 10.0 software.

### Preparation and staining of cell suspensions for flow cytometry

Ear skin samples were split in half, cut into small pieces and digested in 1ml of RPMI-1640 medium with 198U/ml DNase I (#10104159001), 0.03125mg/ml LiberaseTM (#5401119001) and 0.25mg/ml hyaluronidase (#H3506, all Sigma) at 37°C and 1400rpm for 1h and filtered (100µm strainer). For the analysis of cells in the blood, the thoracic cavity was opened using surgical scissors and forceps and subsequent puncture of the left ventricle using a 27G needle attached to a 1ml syringe. 80µl blood was mixed with 40µl heparin (#H3149, Sigma; 500U/ml), erythrocyte lysis was performed in 4ml lysis buffer (155mM NH_4_Cl, 10mM KHCO_3_, 0.13mM EDTA in PBS) for 4min and samples were washed with PBS. Inguinal lymph nodes were dissociated by the use of 2 object slides. Single cell suspensions were incubated with an anti-FcγRIII/II antibody (1:100, #101320, BioLegend) in PBS/2mM EDTA/0.5% BSA for 15min at 4°C. Cells were stained with primary mAbs in 100µl for 15min at 4°C, washed and fixed using PBS/0.5% formaldehyde. For some samples, intracellular staining was performed (#88-8824-00, ThermoFisher), according to manufacturer’s instructions. If reporter MCs were analyzed, no fixation was performed. Samples were acquired with a LSR Fortessa I or LSR Fortessa II flow cytometer (BD, San Jose, CA) and analyzed with FlowJo software (Ashland, OR, USA).

### Generation of peritoneal cell-derived mast cells (PCMCs)

Peritoneal cell-derived mast cells (PCMCs) were generated as previously described (67). Briefly, peritoneal cavity was rinsed with 5ml PBS and cells were incubated afterwards in RPMI 1640 with 10% FCS, 100µg/ml penicillin/streptomycin, 50µM β-mercaptoethanol, 1mM sodium pyruvate, 10ng/ml IL-3, and 30ng/ml SCF. Non-adherent cells were carefully removed after 48h and fresh culture medium was supplied. Starting from d5, MC culture medium (supplemented with 10ng/ml IL-3 and 10ng/ml SCF) was changed twice a week. Enriched PCMCs were used at d8-d14 for further analysis.

### PCMC degranulation assays and cytokine measurement

2×10^5^ PCMCs were seeded in either MC culture medium only or supplemented with 300ng/ml anti-DNP IgE (#D8406, Sigma-Aldrich) for sensitization. After 24h, for some experiments, PCMCs were treated with a PI3K inhibitor LY294002 (#440202, Sigma-Aldrich; 50µM in DMSO) or equal amounts of DMSO for 30min. Then, sensitized PCMCs were stimulated with 100ng/ml DNP-BSA (#A23018, ThermoFisher) and untreated PCMCs with 500ng/ml calcium ionophore A23187 (#C7522, Sigma-Aldrich), 500µM ATP (#10519987001, Roche) or 50ng/ml IL-33 (#210-33, PeproTech) in a total volume of 125µl MC culture medium without phenol red, respectively. After 10min, cell culture plate was placed on ice for 5min and centrifuged (365g, 4°C). Afterwards, cells were stained for flow cytometry or analyzed by β-hexosaminidase. For the latter, 120µl medium was removed, before 125µl lysis buffer (1% triton X-100 in RPMI-1640) was added for 5min. Supernatants and lysates were stored at –80°C. β-hexosaminidase assay was performed as described before (71). Briefly, 25µl of each supernatant and lysate were mixed with 25µl 4mM pNAG solution (4-nitrophenyl-N-acetyl-β-D-glucosaminid (#N9376, Sigma-Aldrich) in citric acid), incubated for 1h at 37°C and 150µl 200mM glycine in PBS (pH 10.7) was added. Absorption at 405nm and 630nm was measured with a microplate reader with automatic background subtraction. Percentage of β-hexosaminidase release was calculated as ratio of absorption of supernatant divided by the sum of absorption of supernatant and lysate. Cytokine concentrations were measured using a customized LegendPLEX, according to the manufacturer’s instructions. Therefore, PCMCs were stimulated as described above. After incubation for 6h at 37°C, supernatants were carefully collected and stored at –80°C before used for measurement.

### Live cell imaging of PCMC degranulation

1×10^5^ IgE-sensitized (300ng/ml anti-DNP-IgE (#D8406, Sigma), overnight) or non-sensitized PCMCs were placed in a poly-L-lysine coated (0.01% in ddH_2_O, #P8920, Sigma) 18 well µ-slide (#81811, ibidi) in Tyrode’s buffer (130mM NaCl, 5mM KCl, 1.4mM CaCl_2_, 1mM MgCl_2_, 10mM HEPES, 5.6mM Glucose, 0.1% BSA in ddH_2_O, pH 7.4), supplemented with 5µg/ml avidine-sulforodamine 101 (Av.SRho, #A2348, Sigma) and 1x CellBrite Steady 405 (#30105-T, Biotium). Cells were allowed to adhere for 30min, before stimulus (1ng/ml DNP-BSA (#A23018, ThermoFisher) or 5µM ATP (#10519987001, Sigma)) was added. Immediately following the configuration of all requisite parameters, the image acquisition process was initiated 1min after stimulation. Technical setup was an inverted confocal laser scanning microscope TCS SP8 (Leica Microsystems, Wetzlar, Germany) with stage-top incubation chamber with 37°C and 5% humidified CO_2_. Illumination was performed via an x63 oil dipping lens with 1.4 NA and a 12kHz resonant scanner (Leica Microsystems, Wetzlar, Germany). Images were recorded as 12-bit z-stacks with a resolution of 512×512 pixels (scaling: 92×92µm) as a z-stack of 20µm (z spacing: 0.4µm) every 10.4s. For visualization and quantification, 4 to 8 images per sample were acquired of randomly chosen ROI. Image analysis was performed using FIJI software. Briefly, a median filter was applied before creating a maximum intensity projection of the entire z-stack. Mean intensities were measured for selected cells over the time of image acquisition using the measurement tool.

### Measurement of intracellular Ca^2+^ levels

Intracellular Ca^2+^ levels were analyzed by ratiometric measurement of Fura-2 emission at 510nm after excitation at 340/380nm at a Tecan Infinite plate reader. 2×10^5^ PCMCs were loaded with 2µg/ml Fura-2 (#F1221, ThermoFisher) in MC culture medium for 30min, washed and seeded into black 96-well plates with translucent bottom, coated with 0.01% poly-L-lysine (#P4707, Sigma), for 15min. After washing, PCMCs were resuspended in either 0mM or 2mM Ringer’s solution (155mM NaCl, 4.5mM KCl, 3mM MgCl_2_, 5mM HEPES, 10mM Glucose, 0mM or 2mM CaCl_2_ in ddH_2_O, pH 7.4). When resuspended in 2mM Ringer’s solution, signal was acquired for 600s after stimulation with 500µM ATP (#10519987001, Roche). Otherwise, when resuspended in 0mM Ringer’s solution, baseline signal was acquired for 135s before Ca^2+^ store depletion was induced by stimulation with 500µM ATP. After additional 485s, PCMCs were supplemented with an equal volume of 2mM Ca^2+^ Ringer’s solution (final extracellular Ca^2+^ concentration: 1mM) and measurement was performed for a total of 18min. Store depletion and SOCE were calculated as highest peak and area under the curve in the respective segments. Baseline Ca^2+^ influx was quantified as non-normalized Fura-2 (340/380nm) signal until ATP was added.

### Western blot analysis

For cell lysis, lysis buffer containing 20mM HEPES, 10mM EGTA, 40mM β-glycerophosphate, 2.5mM MgCl_2_, 2mM orthovanadate, 1mM dithiothreitol, 20μg/mL aprotinin, 20μg/mL leupeptin and 1% Triton X-100 (pH 7.5) was used. The protein content was determined by using the BCA Protein Assay Kit (Pierce). Afterwards, samples were boiled in Laemmli buffer. The samples were separated in 10% sodium dodecyl sulphate (SDS)-Laemmli gels and transferred onto nitrocellulose membranes (SERVA) by western blotting. Membranes were subsequently blocked with dry milk, washed (0.1% Tween 20 in TBS) and incubated with primary antibodies overnight. We used pSHIP1 (#3941), pAKT (#9271) and HSP90 (#4874) (all from cell signaling technology). Membranes were washed (0.1% Tween 20 in TBS) and incubated (4h) with the HRP-conjugated anti-rabbit-Ig antibody (#5220–0336, Medac GmbH). For protein detection, the ECL Western Blotting Substrate (Pierce) was used.

### Statistical analysis

Analyses were performed using GraphPad Prism software (version 10.5.0). Unless reported otherwise, data is shown as mean±SEM (bar graphs, line graphs) or box plots (minimum to maximum) with each dot representing a biological replicate. For big data sets, exclusion methods were applied to identify outliers. Unpaired nonparametric Mann-Whitney U-test was used for the comparison of two groups and non-paired nonparametric Kruskal-Wallis test or mixed-effects analysis for the comparison of more than two groups, respectively. Asterisks indicate significance (*p<0.05, **p<0.01, ***p<0.001, ****p<0.0001, ns – not significant). Flow cytometry and microscopy images are representative of individual experiments.

## SUPPLEMENTARY MATERIALS

Supplementary Figure S1 to S3

Supplementary Video S1 to S3

## ACKNOWLEDGEMENTS

We cordially thank all members of the Dudeck laboratory for discussions and support. Expert technical support and expertise by J. Kotrba, M. Voss, and S. Kliche (Magdeburg, Germany) is gratefully acknowledged. We thank R. Fässler (Munich, Germany) for providing *Itgb1^FL/FL^* mice for breedings. Graphical abstract, icons and schematic representations (except Fig. 3I) were created in BioRender. Hoffmann, A. (2026) https://BioRender.com/jlfc6f0. This work was funded by the Deutsche Forschungsgemeinschaft (DFG, German Research Foundation): Project-ID 361210922/RTG2408/TP4, DU1172/8-1 and DU1172/9-1 to AD.

## AUTHOR CONTRIBUTIONS

AH and AD conceived the project and wrote the manuscript. AH, SD, RI, KKD JD, KB, CK, and TF performed experiments, analyzed and compiled the data. SF, SK, and MS supported the project, interpreted and discussed the data, wrote and edited the manuscript.

## SUPPLEMENTARY FIGURES

**Supplementary Figure S1.**
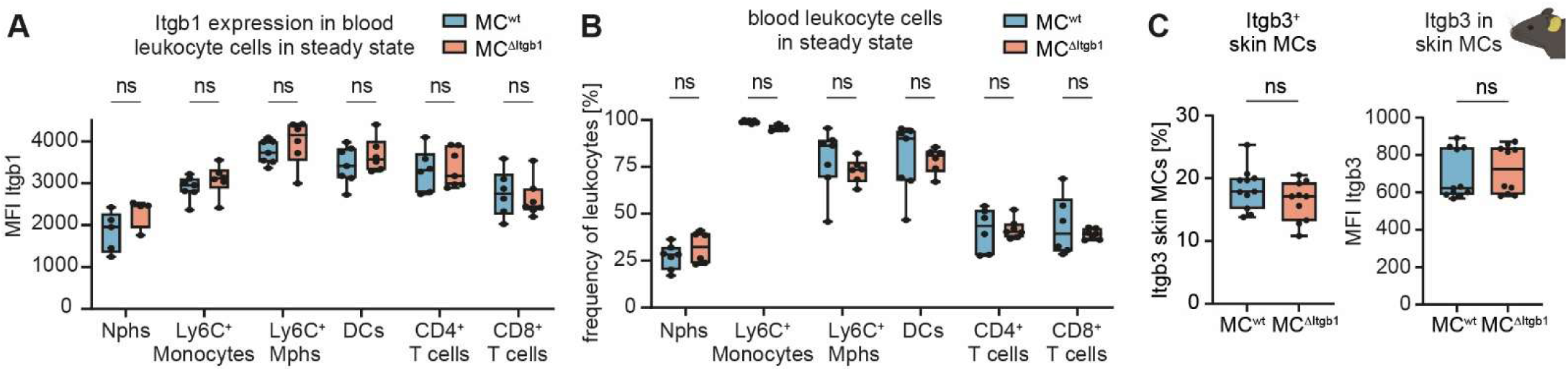
Evaluation of integrin expression in leukocytes verifies specificity of the conditional Itgb1 knock-out in MCs. (A) Expression of Itgb1 as median fluorescence intensity (MFI) and (B) frequency of blood leukocyte subsets in steady state (n=5-7 (MC^wt^) and n=4-7 (MC^ΔItgb1^). (C) Frequency of ear skin MCs and (D) expression level of integrin β3 (Itgb3) as MFI (n=11 (MC^wt^) and n=10 (MC^ΔItgb1^). DCs – dendritic cells, Mono – monocytes, Mphs – macrophages, Nphs – neutrophils. ns – not significant.

**Supplementary Figure S2.**
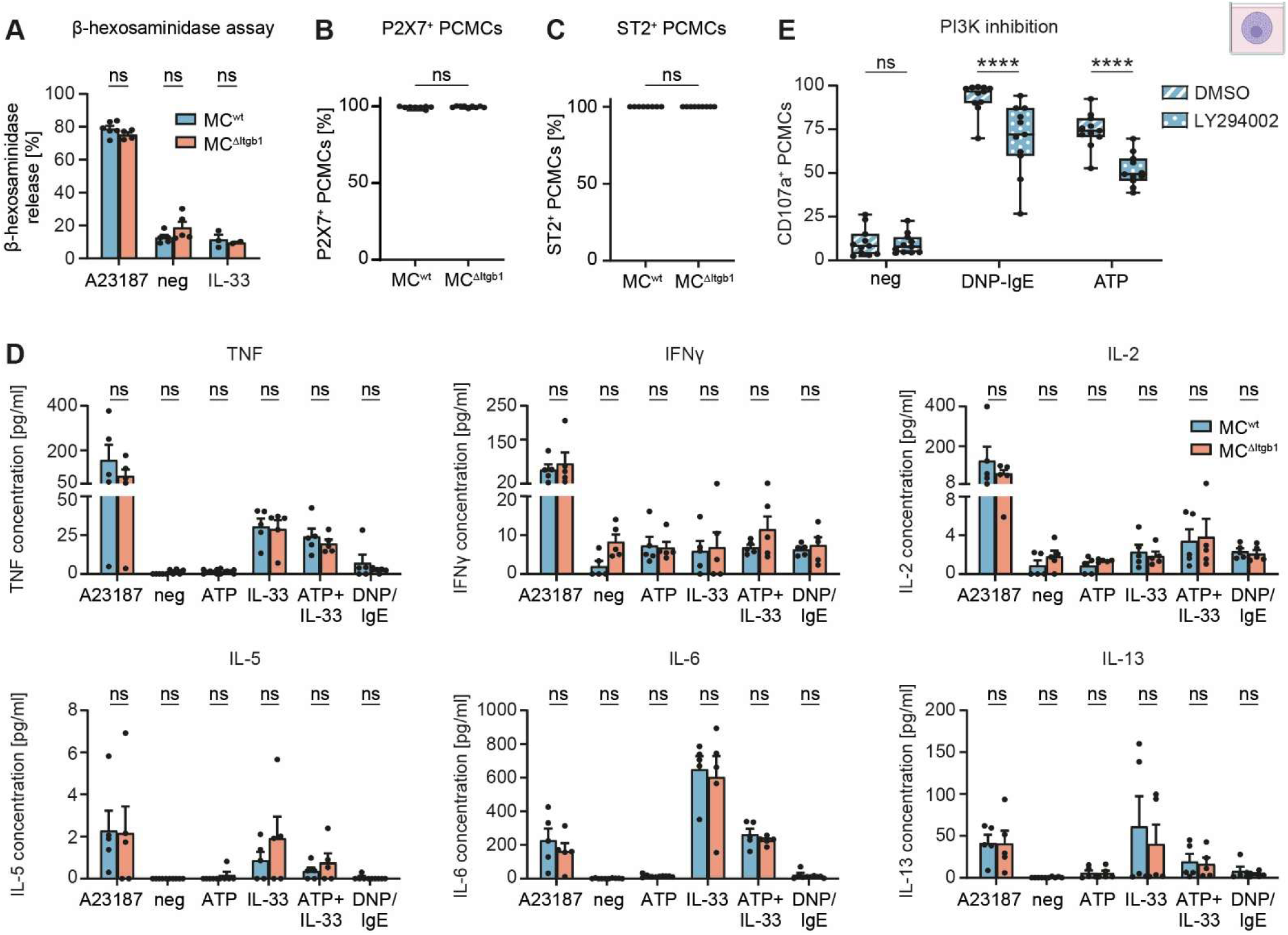
Assessment of degranulation and receptor expression of PCMCs *in vitro*. (A) β-hexosaminidase release assay upon stimulation with calcium ionophore A23187 (positive control), medium (negative control) or 50ng/ml IL-33 (n=4-6 (MC^wt^) and n=4-5 (MC^ΔItgb1^). Frequency of (B) P2X7^+^ and (C) ST2^+^ PCMCs (n=8 (MC^wt^) and n=9 (MC^ΔItgb1^). (D) TNF, IFNγ, IL-2, IL-5, IL-6 and IL-13 concentration in the supernatant upon stimulation of PCMCs with A23187, medium, 500µM ATP, 50ng/ml IL-33, 500µM ATP + 50ng/ml IL-33 or 100ng/ml DNP-BSA (sensitized with anti-DNP IgE), respectively (n=5). (E) Inhibition of MC degranulation upon stimulation with 100ng/ml DNP-BSA (sensitized with anti-DNP IgE), 500µM ATP or medium (negative control), after treatment with PI3K inhibitor LY294002 or DMSO for 30min, respectively (n=10 (MC^wt^) and n=12 (MC^ΔItgb1^). **** p<0.0001, ns – not significant.

**Supplementary Figure S3.**
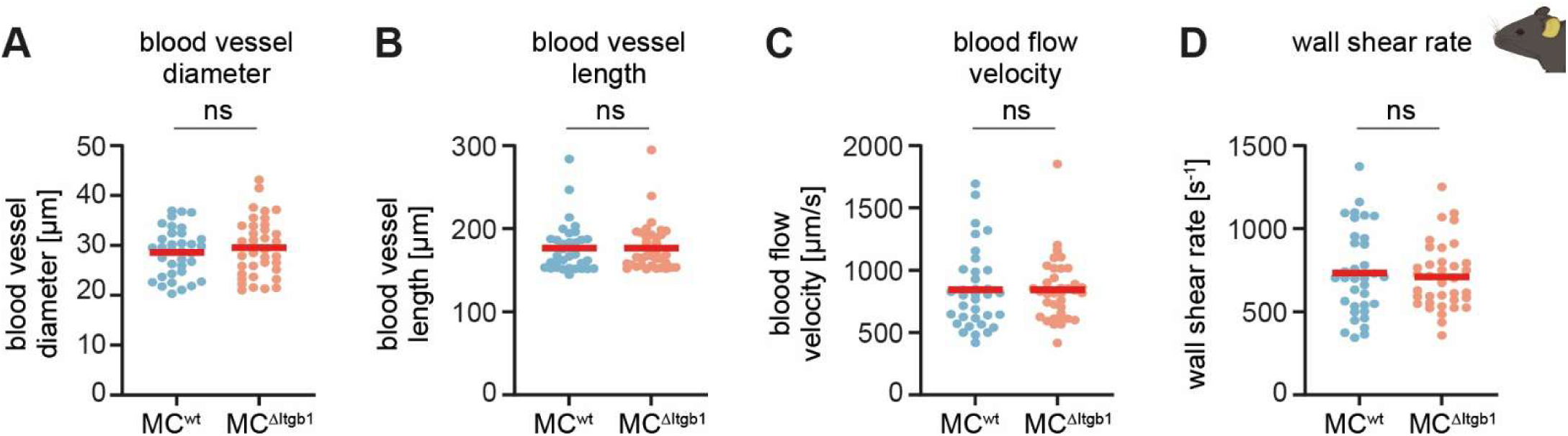
Hemodynamic parameters of analyzed blood vessels are similar during intravital 2-photon microscopy analysis. (A) Blood vessel diameter, (B) blood vessel length, (C) blood flow velocity and (D) wall shear rate of analyzed blood vessels during intravital 2-photon microscopy (n=35 (MC^wt^) and n=37 (MC^ΔItgb1^) vessels of N=5 mice per group). Red bars indicate mean; ns – not significant.

**Supplementary Video S1. Live-cell imaging of MC degranulation upon ATP stimulation *in vitro*.** Representative video of *in vitro* live-cell confocal laser scanning microscopy of MC^wt^ and MC^ΔItgb1^ upon stimulation with 5µM ATP for 30min. MC degranulation is visualized by binding of avidin sulforhodamine 101 to heparin in MC granules, thereby labeling exocytosed MC granules. Following the membrane disintegration during MC degranulation, intracellular granules will also be stained by avidin. Green – CellBrite steady 405, red – avidin sulforhodamine 101, visualization as maximum intensity projection (MIP) of single channels (left and middle) and merge (right). Scale bar 5µm. See also Fig. 4.

**Supplementary Video S2. Live-cell imaging of MC degranulation upon DNP/IgE stimulation *in vitro*.** Representative video of *in vitro* live-cell confocal laser scanning microscopy of MC^wt^ and MC^ΔItgb1^ upon stimulation with 1ng DNP-BSA of anti-DNP IgE sensitized PCMCs for 30min. MC degranulation is visualized by binding of avidin sulforhodamine 101 to heparin in MC granules, thereby labeling exocytosed MC granules. Following the membrane disintegration during MC degranulation, intracellular granules will also be stained by avidin. Green – CellBrite steady 405, red – avidin sulforhodamine 101, visualization as maximum intensity projection (MIP) of single channels (left and middle) and merge (right). Scale bar 5µm. See also Fig. 4.

**Supplementary Video S3. Intravital 2-photon microscopy of intraluminal neutrophil rolling and adhesion upon CHS.** Representative video of intravital 2-photon microscopy of intraluminal neutrophil rolling in the ear skin of MC^wt^ and MC^ΔItgb1^ mice 12h after epicutaneous application of 1-fluoro-2,4-dinitrobenzene (DNFB). Rolling and adhesion of individual neutrophils was analyzed as previously described (69) for 1min. Red – CD31-AF647, cyan – Ly6G-PE. Scale bar 20µm. See also Fig. 6.

